# A conserved, non-canonical insert in FIS1 is required for mitochondrial fission and recruitment of DRP1 and TBC1D15

**DOI:** 10.1101/2023.01.24.525364

**Authors:** UK Ihenacho, R Toro, RH Mansour, RB Hill

**Affiliations:** Department of Biochemistry, Medical College of Wisconsin, Milwaukee, Wisconsin, USA

**Keywords:** Mitochondria, mitophagy, peroxisome, nuclear magnetic resonance (NMR), fission, tetratricopeptide repeat, repeat proteins, protein motif, organelle dynamics, dynamin

## Abstract

Mitochondrial fission protein 1 (FIS1) is conserved in all eukaryotes yet its activity in metazoans is thought divergent from lower eukaryotes like fungi. To address this discrepancy, structure-based sequence alignments revealed a conserved but non-canonical, three-residue insert in a turn of FIS1, suggesting conserved activity. In vertebrate FIS1 this insert is serine (S45), lysine (K46), and tyrosine (Y47). To determine the biological role of this “SKY insert”, three variants were evaluated for their fold, and tested in HCT116 cells for altered mitochondrial morphology and recruitment of effectors, DRP1 and TBC1D15. Substitution of the SKY insert with three alanine residues (AAA), or deletion of the insert (ΔSKY), did not substantially alter the fold or thermal stability of the protein. Replacing SKY with a canonical turn (ΔSKYD49G) introduced significant conformational heterogeneity by NMR that was removed upon deletion of a known regulatory region, the FIS1 arm. Expression of AAA fragmented mitochondria into perinuclear clumps associated with increased mitochondrial DRP1 similar to the wild-type protein. In contrast, expression of ΔSKY variants elongated mitochondrial networks and reduced mitochondrial DRP1. Co-expression of YFP-TBC1D15 partially rescued mitochondrial morphology and DRP1 recruitment for ΔSKY variants, although ΔSKY variants were markedly unable to support TBC1D15 assembly into punctate structures found upon co-expression with wildtype FIS1 or the AAA variant. Collectively these results show that FIS1 activity can be modulated by conserved residues supporting a generalized model whereby FIS1 is governed by intramolecular interactions between the regulatory FIS1 arm and SKY insert that may be conserved across species.

## Introduction

In most eukaryotes, mitochondria exist as highly dynamic networks that balance frequent fission and fusion events to maintain the appropriate morphology necessary for organelle function and cellular homeostasis (1–4). Early gene complementation screens in yeast revealed model genes, Dnm1, Mdv1/Caf4, and FIS1, involved in mitochondrial fission (5–10). These experiments led to a proposed rudimentary fission apparatus comprised of a resident outer-membrane protein (Fis1p) acting in concert with an adaptor (Mdv1p/Caf4p) to recruit a GTPase mechanoenzyme (Dnm1p) from the cytoplasm to sites of scission (8, 9, 11–14). However, only FIS1 and DRP1 are present in all mitochondria-bearing species, and Mdv1/Caf4 are fungal-specific with no known vertebrate orthologs identified to date. Moreover, increasing FIS1 expression potently induces DRP1-dependent division of target organelles – mitochondria (15–17), peroxisomes, and plastids – regardless of species (18–22). These considerations suggest that FIS1 and DRP1 constitute the core fission machinery, and that adaptors are unique from species to species. Supporting this idea are sequence alignments of FIS1 and DRP1 that show high conservation across species (23, 24). Despite these considerations, the fission machinery in vertebrates is more complex as additional mitochondrial proteins like MFF and MID49/51 potently recruit DRP1 in the absence of FIS1(25–29). Groundbreaking work also identified that FIS1 recruits mitophagy adapters TBC1D15/17 to mitochondria suggesting additional roles for FIS1 in vertebrates (30–32). In support, studies from simple to complex eukaryotes have described specific roles for FIS1 in peripheral mitochondrial fission during stress and/or development (33), with recent super-resolution microscopy revealing that MFF recruits DRP1 for mid-body, housekeeping fission, whereas FIS1 recruits DRP1 to peripheral sites for mitophagic removal (34). FIS1’s transmembrane and TPR domains are indispensable for the mitochondrial recruitment of TBC1D15, although, the exact mechanisms are not completely understood (30). These findings suggest that FIS1 functional mechanisms have diverged in vertebrates despite the amino acid sequence conservation, thus raising the question of what governs FIS1 recruitment of TBC1D15/17 or DRP1.

Insights into FIS1 activity may be gained by consideration of its structure, which has two domains: a C-terminal transmembrane domain that anchors it to membranes and a soluble helical domain that adopts a fold reminiscent of tetratricopeptide repeat (TPR) proteins (**Figure 1A**) (35, 36). TPRs are 34 amino acid degenerate sequences that form a helix-turn-helix motif occurring as three or more repeats to form superhelical arrays. This architecture creates a concave and convex face that mediates binding to multiple partners (37). To date, most TPRs seem to mediate binding via their concave face, access to which is often regulated by steric occlusion from flanking regions (38). FIS1 is an atypical TPR protein because it possesses two repeats, only one of which is canonical (35, 36). Furthermore, FIS1 exists in oligomeric heterocomplexes mediated by its TPRs, which may be auto-inhibited by its N-terminal helix as deletion of this helix enhances FIS1 oligomerization and DRP1 recruitment (17, 23, 39–41). Adjacent to the N-terminal helix is a disordered region of FIS1, termed the FIS1 arm, that is required for its mitochondrial fission functions in both yeast and human cells (41, 42). Consistent with a key role for the N-terminal region are splice variants in mice and fruitflies that lack this region (43).

**Figure 1.**
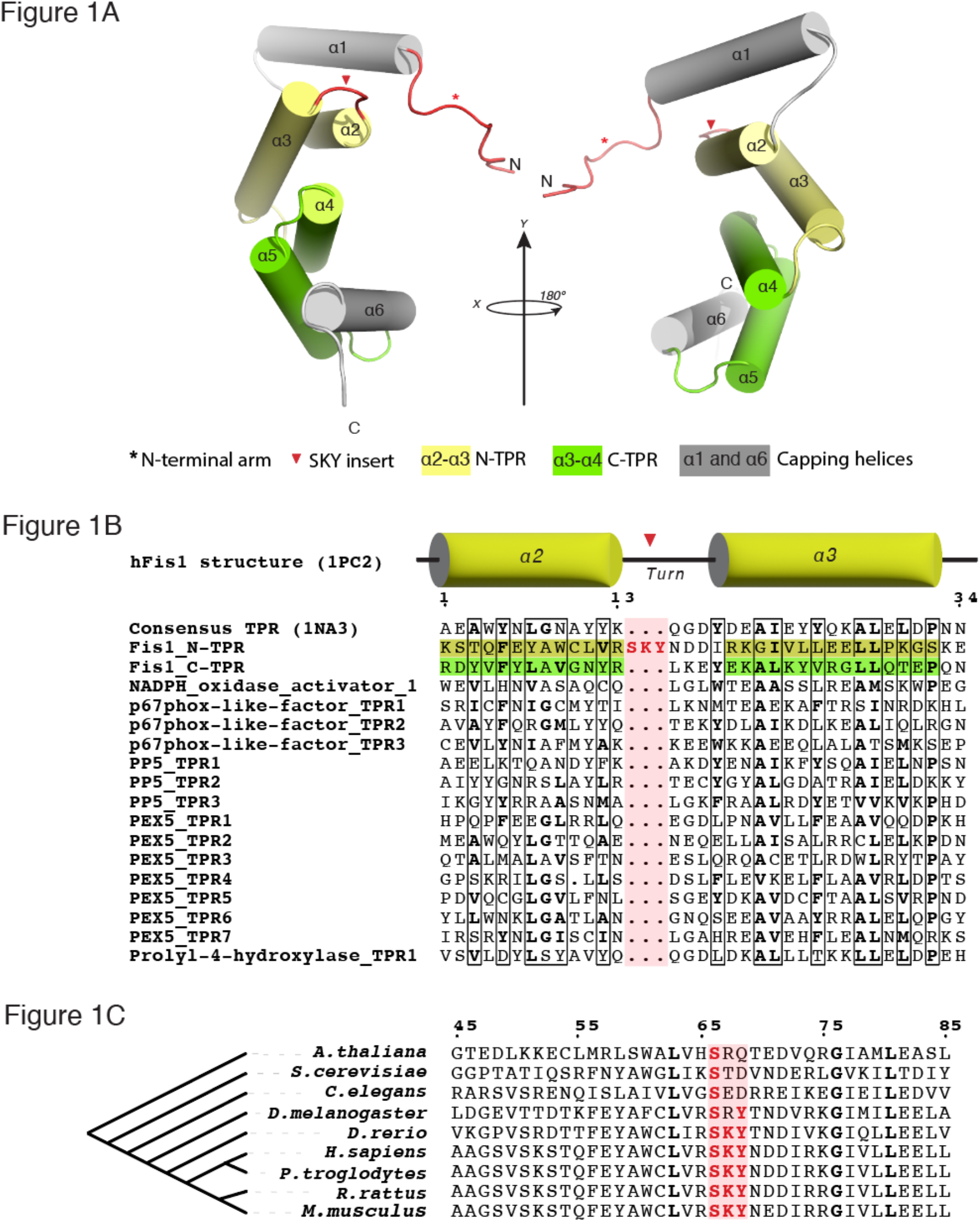
Structure-based sequence alignments reveal a conserved three-residue insert in the N-terminal TPR of FIS1. **A,** Solution structure of human FIS1 (PDB: 1PC2) depicting the N-terminal region called the “arm” (Red asterisk), two tetratricopeptide repeats; the N-TPR in yellow (α-helices 2-3), and the C-TPR in green (α-helices 4-5) with flanking α-helices 1 and 6 in gray. The SKY insert (red arrowhead) is found in the turn of N-TPR in red between α-helices 2 and 3. **B**, Structure-based sequence alignments of human FIS1’s tetratricopeptide repeats and TPRs in the human proteome. The six-helix consensus TPR protein structure (PDB:1NA0) was used as a template. Note that three-residues (Ser45, Lys46, Tyr47 in human) are inserted in the canonical TPR turn. **C,** The three-residue insert is conserved across FIS1 species and is always SKY in vertebrates.

In the current study, we searched for unifying mechanisms that could account for conservations of FIS1 functions and the observed differences across species. Structure and phylogeny-based sequence alignments revealed a three-residue insert in the N-terminal TPR that is uniquely conserved in all species. Moreover, this insert is conserved as Ser-Lys-Tyr (SKY) in all vertebrates. Here we report that the conserved SKY insert is not a stringent structural requirement for human FIS1 but is indispensable for its mitochondrial recruitment of TBC1D15 complexes that appear crucial to FIS1’s mitochondria division functions in vertebrates. Furthermore, we show that FIS1-induced fission of mitochondria networks can be potently up or downregulated by simply perturbing insert residues. Overall, our findings provide useful insights into elucidating unifying structural mechanisms that govern FIS1 functions in eukaryotes.

## Results

### FIS1 has a conserved three-residue insert in the first TPR

We used structure-based sequence alignments to compare human proteins containing TPRs with FIS1 (**Figure 1B**). Strikingly, these alignments revealed a non-canonical TPR feature in the first, but not the second, TPR of FIS1: Instead of the canonical 34-amino acids that define a TPR, FIS1’s first TPR (N-TPR) contains an additional stretch of three amino acids – serine, lysine, and tyrosine – inserted within the turn region of the canonical helix-turn-helix of a TPR (**Figure 1B**). Curiously, a three-residue insert is present in all known FIS1 sequences and occurs as an invariant SKY in vertebrates (**Figure 1C**). As the “SKY insert” is not required to specify the TPR fold, we infer that it is not conserved for structural purposes, but rather for FIS1 activity.

### Rational design and validation of SKY variants

To investigate the functional relevance of the SKY insert, we designed a FIS1 variant with a short canonical TPR turn lacking the insert. This was accomplished by analyzing the TPRs from a well-characterized consensus TPR sequence that adopts the canonical structure. This consensus TPR is an entirely non-native sequence designed from statistical thermodynamic analysis of TPR sequences and was shown to fold into the desired TPR structure indicating the robustness of the design and TPR fold. Structural comparison of the FIS1 N-TPR with the consensus TPR from CTPR3 (1NA0.pdb) showed excellent alignment of the two helices (Ca RMSD = 1.1Å) with only a slightly longer turn for FIS1 (**Figure 2A**). This suggested that replacing the SKY insert with the turn from the TPR consensus sequence would not perturb the protein fold. TPRs have a characteristic three-residue turn with φ, ψ backbone torsional angles that, according to Effimov’s convention (44), correspond to γ-α_L_-β of Ramachandran space with the central residue typically, but not always, being a GLY that can readily adopt α_L_ values of φ, ψ space. Commonly the third position is a small, hydrophilic residue that adopts β space. Consistent with these principles, the consensus TPR turn is specified by the sequence Q-G-D, whereas the FIS1 turn is S-K-Y-N-D-D, with the SKY insert occurring before position 1. Deletion of the SKY insert leaves N-D-D to serve as the turn, which compares favorably to the consensus turn residues Q-G-D with the exception of the central GLY. Based on these considerations, we made four constructs by (i) substituting three Ala residues for SKY (AAA), (ii) deleting the SKY insert (ΔSKY) that retains the central Asp to give N-D-D, (iii) deleting the SKY insert that substitutes the central Asp (D49) with the canonical Gly (ΔSKYD49G) to give N-G-D, and (iv) a control that retained the SKY insert but replaced the succeeding Asp with Gly (D49G).

**Figure 2.**
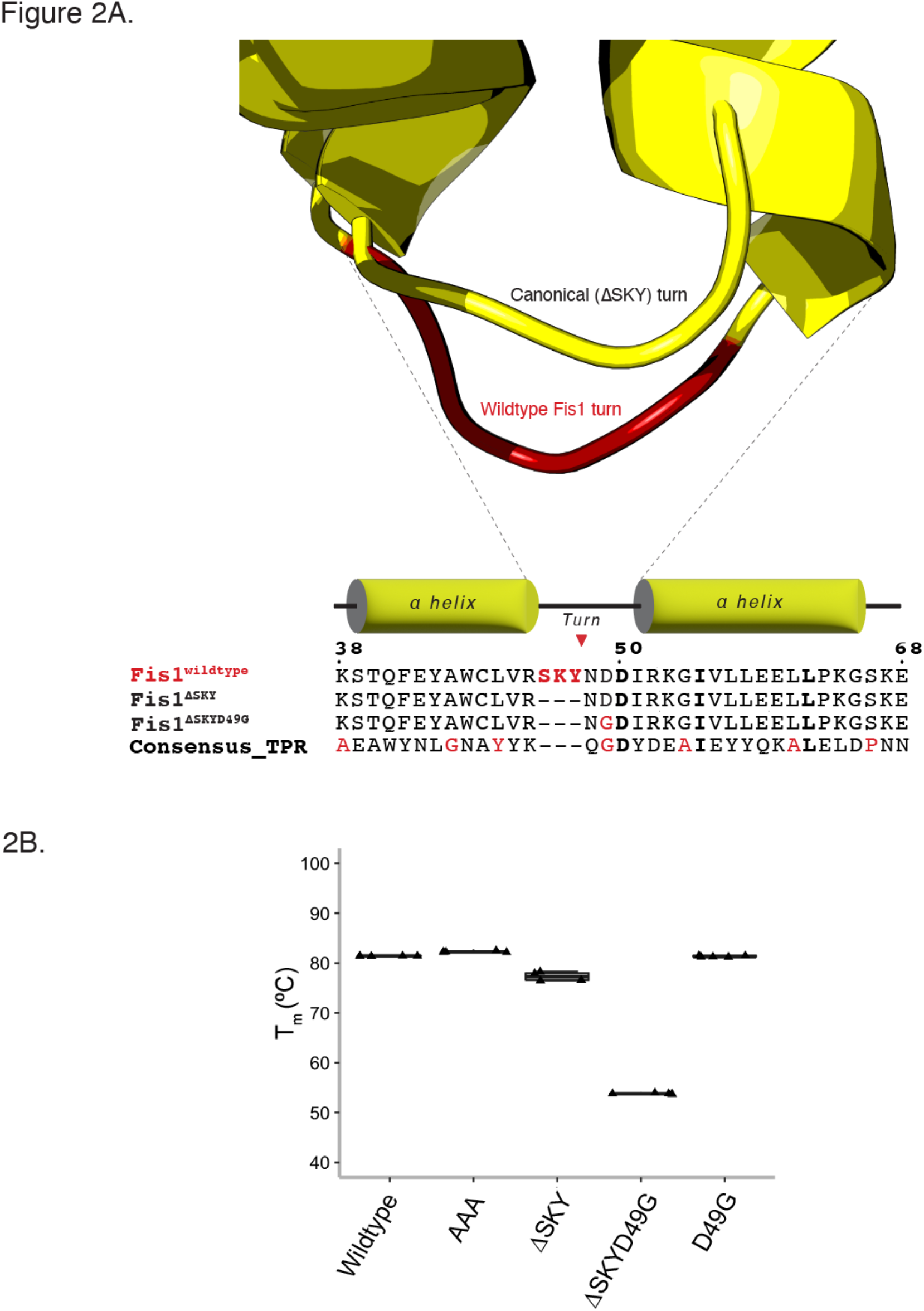
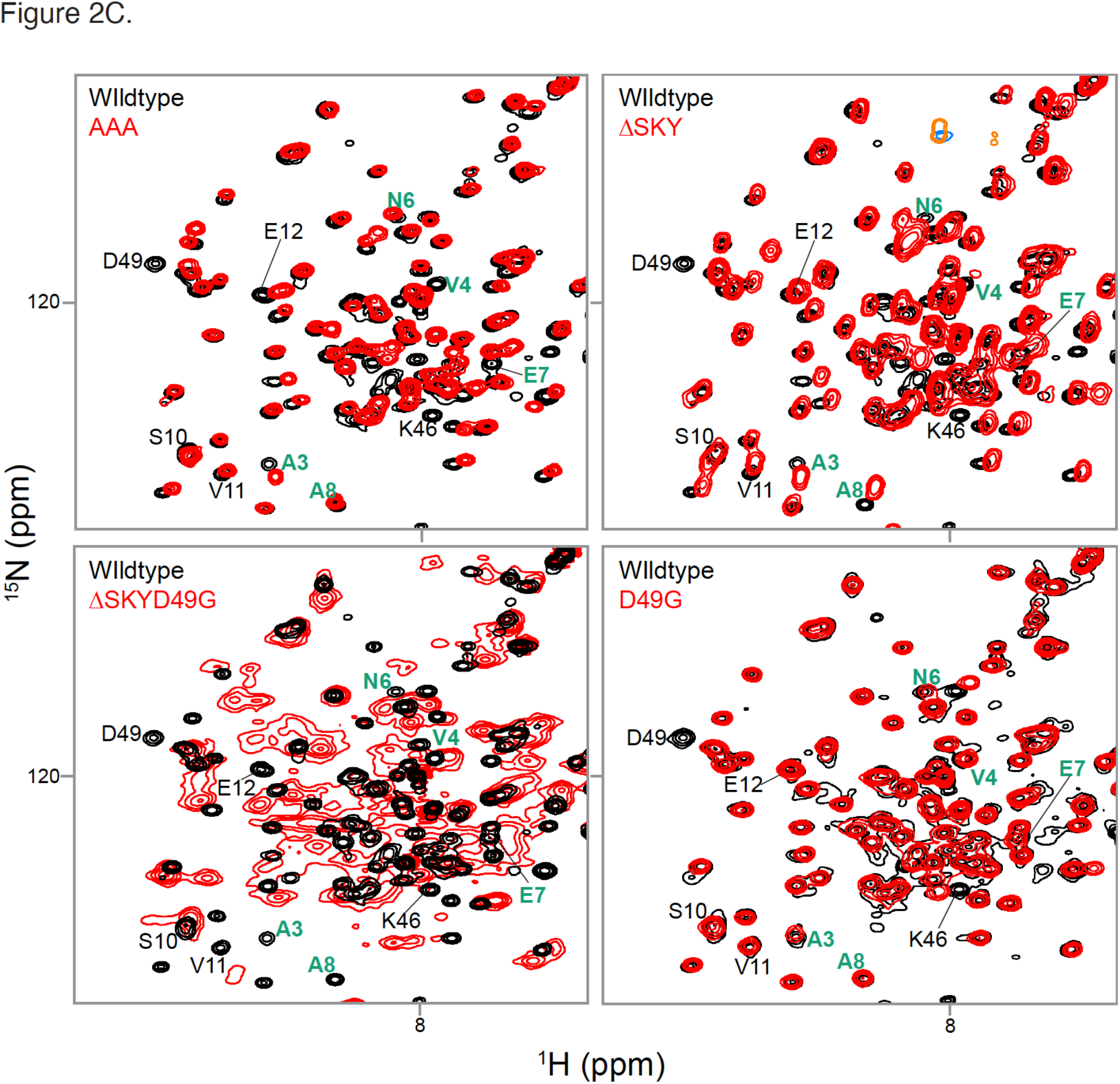
Rational design and validation of a ΔSKY FIS1 variant. **A**, Superposition of the N-TPR turn of FIS1 (PDB: 1PC2) with a canonical TPR turn from the rationally-designed, consensus TPR protein (PDB:1NA0). The ΔSKY construct removes the insert and ΔSKYD49G substitutes with a conserved Gly, see text for rationale. **B**, The midpoint of the thermal unfolding transition was determined by fitting light scattering data collected from 25-95°C with the mean ± standard deviation from 3-5 technical replicates shown as a box-and-whisker plot. **C**, ^1^H-^15^N HSQC spectral overlays of FIS1 wildtype (black) with indicated variants (red). Data were collected on 100 μM samples at 25 °C, pH 7.4 at 14.1 T. FIS1 arm crosspeaks are indicated in green. See **FigureS1** for full spectra overlays.

To assess the quality of our designs, we recombinantly expressed and purified the cytoplasmic domain of these proteins for biophysical characterization. Thermal stability was measured by monitoring the intrinsic fluorescence with increasing temperature to determine the midpoint of the unfolding transition (T_m_). The wildtype cytoplasmic domain is quite thermally stable with a T_m_ of 81.8 ± 0.1 °C and neither alanine substitutions (AAA) nor the control construct (D49G) impacted thermal stability compared to wildtype (**Figure 2B**). Deletion of the SKY insert (ΔSKY) modestly decreased the T_m_ to 71.5 ± 0.2 °C consistent with the assumption that these residues are dispensable for the TPR fold. However, the ΔSKYD49G construct dramatically decreased the T_m_ to 59.9 ± 0.2 °C. To understand this, we turned to two-dimensional NMR spectroscopy of these proteins uniformly labeled with ^15^N that allows for individual residue contributions to the overall protein fold. All constructs showed similar chemical shift dispersion to wildtype indicating well-folded proteins (**Figure 2C**). However, ΔSKYD49G NMR data showed an increased broadening of resonances throughout the spectrum consistent with a significant degree of conformational heterogeneity. Moreover, cross peaks for amino acid residues corresponding to the N-terminal arm were not detected. To test the role of the FIS1 arm in this conformational heterogeneity, we created a ΔSKYD49G variant lacking the N-terminal arm (ΔNΔSKYD49G) and assessed its structure by thermal melt and NMR. Deletion of the FIS1 arm restored the T_m_ to a value similar to ΔSKY (73.6 ± 0.5 °C) and showed largely resonances similar to wild type with little indication of conformational heterogeneity (**Figure S1**). We interpret these data to indicate that the presence of the N-terminal arm was responsible for inducing conformational heterogeneity in ΔSKYD49G.

### The SKY insert is required for FIS1-induced changes in mitochondrial morphology

To investigate the role of the SKY insert on cellular functions, we transiently expressed wildtype and FIS1 variants along with mitochondrially targeted YFP in human colorectal carcinoma (HCT116) cells. Ectopic overexpression of wildtype FIS1 induces uniform fragmentation and collapse of mitochondrial networks around the nucleus, collectively resulting in perinuclear clumps confirming the findings by others (**Figure 3A**). These changes in morphology were quantified by using MitoGraph to determine the mitochondrial area, which showed a statistically significant decrease for wildtype compared to vector alone (**Figure 3B**). As a control for the TPR domain, a commonly used FIS1 variant (5LA) that replaces five conserved TPR Leu residues with Ala was expressed (17, 30). As previously shown, the 5LA variant also caused mitochondrial clumping with a similar mitochondrial area. Substituting AAA for the SKY insert closely phenocopied wildtype FIS1 with highly fragmented and clumped networks, also with similar mitochondrial areas. By contrast, removal of the SKY insert, in either ΔSKY or ΔSKYD49G, prevented fragmentation and network collapse with an increase of mitochondrial area that was statistically significant. This loss of function was not due to the D49G substitution, which showed mitochondrial morphology similar to wildtype expression.

**Figure 3.**
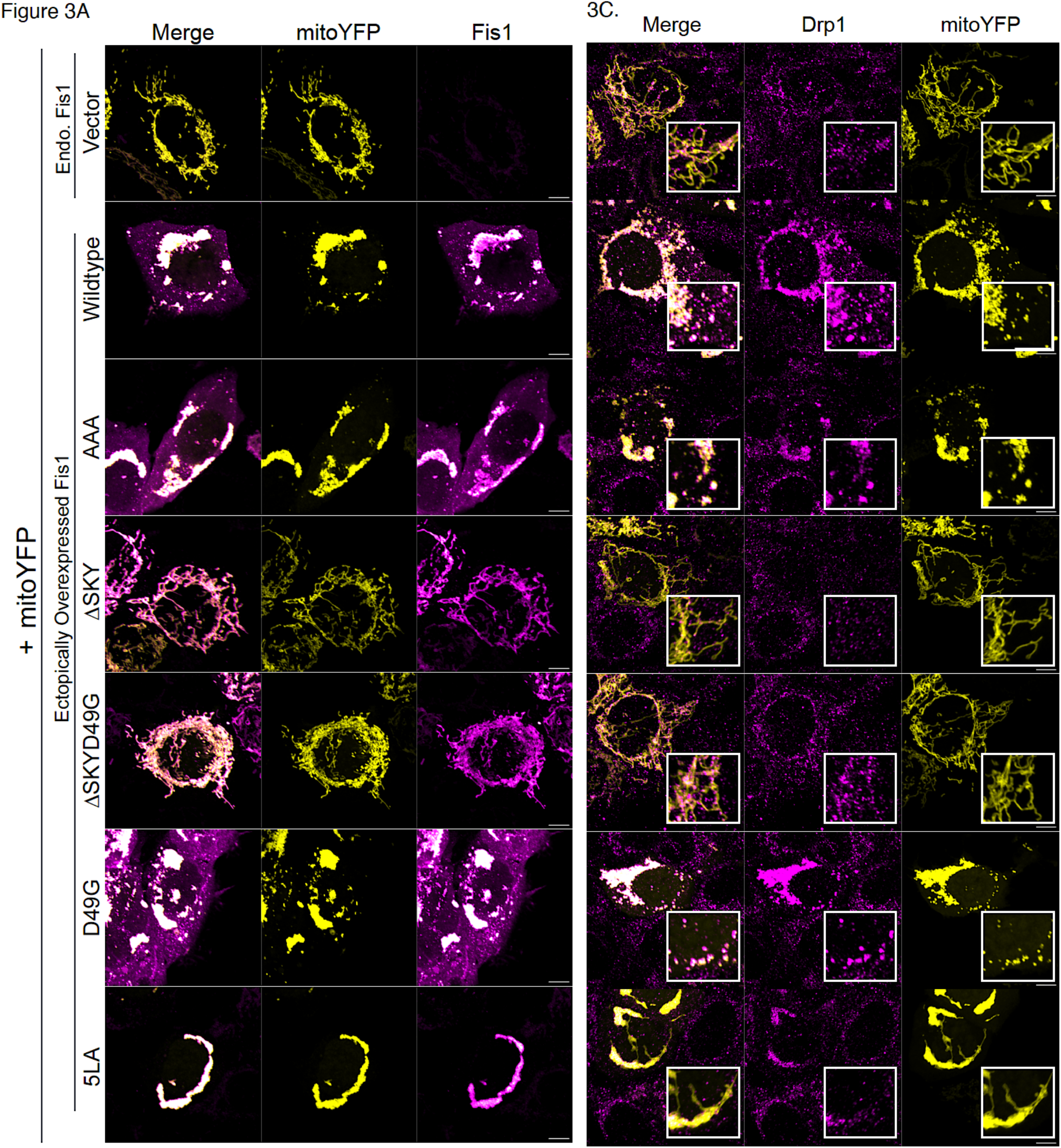
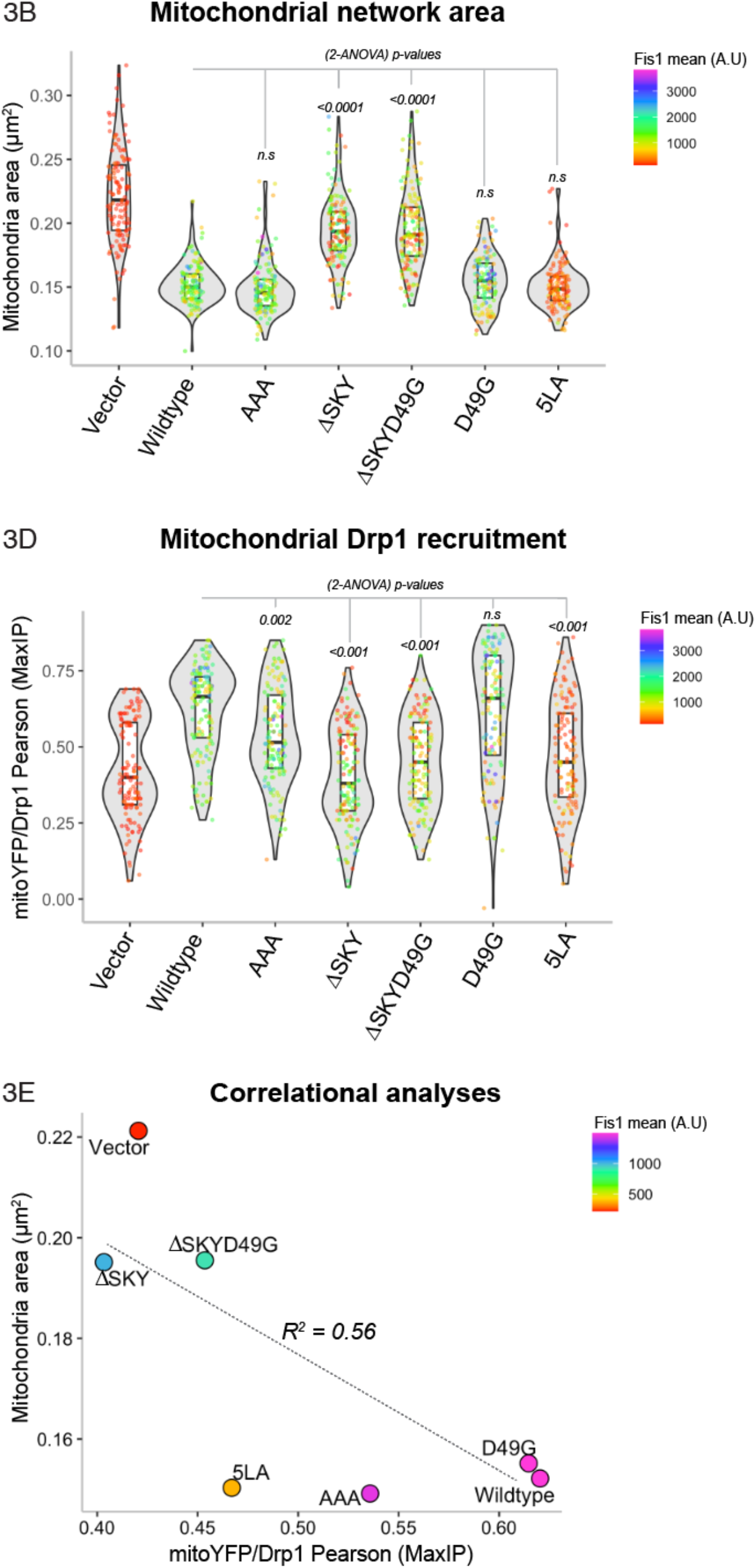
The SKY insert is required for FIS1– induced changes in mitochondrial morphology. HCT116 cells were transfected with mitoYFP and either pcDNA, pcDNA-FIS1 wildtype, or pcDNA-FIS1 variants as indicated, fixed and immunostained sequentially for DRP1, followed by FIS1. **A,** Representative confocal images showing merged anti-FIS1 (magenta-HOT) and mitoYRP (yellow) from single channel images as indicated, Scale bar = 100μm. **B,** Violin plots of average mitochondrial component area. **C,** Representative confocal images showing merged anti-DRP1 (magenta) and mitoYFP (yellow) from single channel images as indicated. Scale bar = 100μm, insets are enlarged 4X with fluorescence intensities adjusted for clarity. **D,** Violin plot of the colocalization between mitoYFP and DRP1 from single cell maximum intensity projections was measured using Pearson’s correlation coefficient. **E,** Correlation plot to determine the relationship between mitochondrial network area and DRP1 recruitment. Each point in **B, D** represents a single cell and each circle in **E** represents the population means, and are colored based on the FIS1 expression levels determined from mean fluorescence intensity per cell. Data represent three biological replicates with *p* values calculated from 2-WAY ANOVA analyses followed by TUKEY honest significant differences (HSD).

The striking morphological changes induced by ectopic FIS1 may involve active mitochondrial recruitment of non-resident factors, such as the highly conserved dynamin family GTPase, DRP1. To evaluate this, we immunostained these cells for DRP1 and quantified colocalization with the MitoYFP signal (**Figure 3C**). Mitochondrial recruitment of DRP1 is potently induced upon ectopic FIS1 overexpression consistent with earlier findings (15, 16, 41) (**Figure 3D**). Therefore, we asked if mitochondrial DRP1 recruitment was significantly perturbed between wild-type and variant FIS1 overexpression. Consistent with an elongated mitochondrial network, mitochondrial DRP1 colocalization decreased by nearly two-fold for both ΔSKY overexpression backgrounds. By contrast, both AAA and 5LA – constructs with clumped networks similar to wildtype – had decreased mitochondrial DRP1 compared to wildtype expression. D49G was able to recruit DRP1 to mitochondria similar to wildtype. The impaired morphology induced by FIS1 variants was reasonably well correlated (R^2^ = 0.56) with their ability to recruit DRP1 (**Figure 3E**), but to a lesser extent for AAA and 5LA indicating that other factors may be involved.

### The FIS1 SKY insert is required for effective mitochondrial recruitment of TBC1D15

The FIS1 TPR domain is exposed to the cytoplasm where it also recruits other binding partners to help govern mitochondrial network morphology. One such class of proteins are the cytoplasmic TBC1 effectors important to many cellular functions including serving as GTPase-activating proteins for Rab family proteins. One TBC1 protein recruited by FIS1 is TBC1D15 and we next asked if the SKY variants impacted TBC1D15 recruitment. For this, the FIS1 constructs were co-transfected with YFP-TBC1D15, and mitochondrial networks were visualized by immunofluorescence to the mitochondrial outer membrane marker TOM20 (**Figure 4A**). In the absence of FIS1 overexpression, the TBC1D15 signal is predominantly cytoplasmic and does not concentrate on mitochondrial networks consistent with endogenous FIS1 levels in HCT116 cells being quite low. By contrast, wildtype FIS1 expression triggers a robust transition of cytosolic TBC1D15 pools onto mitochondrial sites into discrete foci or puncta, which was concomitant with FIS1-induced mitochondrial fragmentation and perinuclear clumping. For FIS1 variants, coexpression of YFP-TBC1D15 impaired the YFP-TBC1D15 cytoplasm-to-puncta transition. To quantify this transition, the mean and mode values of cellular YFP-TBC1D15 signal were measured and reported as mean:mode ratios (**Figure 4B**). For vector alone, the mean and mode are essentially equivalent, reflecting the even distribution of YFP-TBC1D15. For wildtype FIS1 expression, the mean:mode ratio decreases by 40% reflecting a decrease of uniform, cytoplasmic YFP-TBC1D15 and the formation of TBC1D15 punctate structures that reside on mitochondrial surfaces (**Figure 4C**). For AAA, TBC1D15 was recruited into fewer yet larger punctate structures than wildtype as indicated by the increase in mean:mode value. We note that the mitochondrial colocalization values for AAA misleadingly decrease reflecting that a fraction of these large punctate structures are not appropriately captured by our colocalization analysis given their large size, despite being associated with mitochondria. Both ΔSKY constructs showed fewer punctate structures with ΔSKYD49G showing a TBC1D15 recruitment phenotype in between ΔSKY and D49G, the latter of which appeared to recruit more TBC1D15 into punctate structures than wildtype. The 5LA variant, in our hands, impaired formation of TBC1D15 puncta with similar mean:mode ratios to vector alone. We conclude that the SKY insert is also involved in TBC1D15 recruitment.

**Figure 4.**
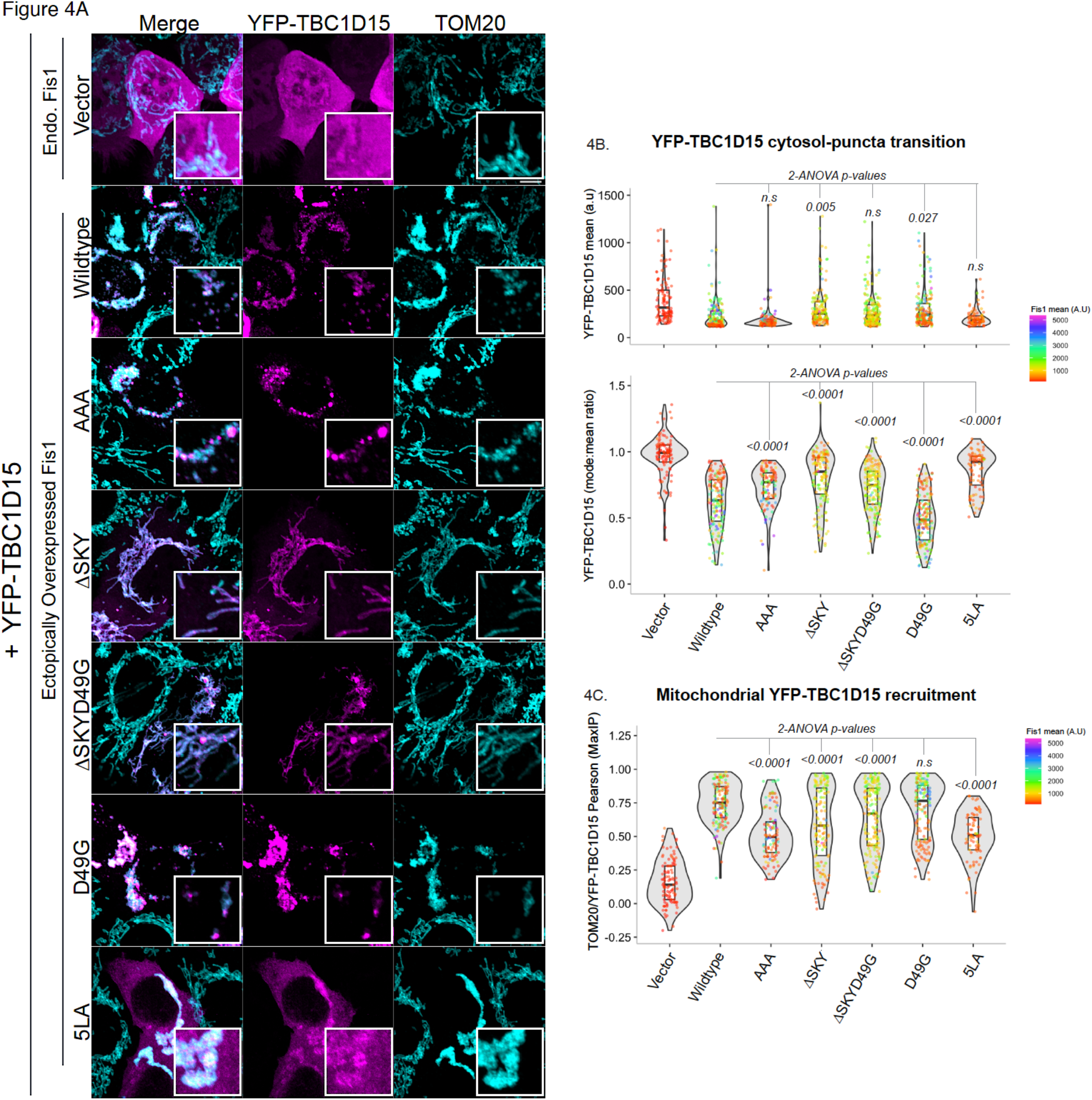
The FIS1 SKY insert is required for effective mitochondrial recruitment of TBC1D15. Analyses of HCT116 cells co-overexpressing FIS1 and YFP-TBC1D15. **A,** From right to left, representative confocal images of cells ectopically expressing YFP-TBC1D15 (magenta) and immunostained TOM20 (cyan) and merges of both channels (merged). Scale bar = 100μm, insets are enlarged 4X. **B,** Violin plots of YFP-TBC1D15 puncta assembly assessed by differences in mode and mean fluorescence intensity values. The top panel shows the mean YFP-TBC1D15 signal intensities, and the bottom panel shows ratios of modal and mean signal intensities. Ratio values close to 1 are indicative of no puncta assembly. **C,** Violin plot of the colocalization between TOM20 and YFP-TBC1D15 from single cell maximum intensity projections was measured using Pearson’s correlation coefficient. Each data point is colored based on the FIS1 expression levels determined from the mean fluorescence intensity per cell. Data represent three biological replicates with *p* values calculated from 2-WAY ANOVA analyses followed by TUKEY honest significant differences (HSD).

### FIS1-mediated fission is potentiated by TBC1D15: loss of DRP1 recruitment is partially rescued by TBC1D15 overexpression

We next asked if co-expression of TBC1D15 with FIS1 variants impacted mitochondrial morphology and DRP1 recruitment. In the absence of exogenous TBC1D15, substitution of the SKY insert with AAA drove a similar clumped morphology to wildtype (**Figure 5A, left panel**), which was quantified again by using MitoGraph to measure the mitochondrial area (**Figure 5B, left panel**). Upon co-expression with YFP-TBC1D15, the mitochondrial area decreased with more resolved mitochondrial clumps (**Figure 5A and B, right panels**) congruent with AAA’s increased YFP-TBC1D15 recruitment (**Figure 4**). Co-expression of either ΔSKY constructs with YFP-TBC1D15 reversed the elongated mitochondrial morphology of these variants as indicated by similar mitochondrial areas to wildtype (**Figure 5A, B**). Co-expression of D49G with YFP-TBC1D15 phenocopied the AAA variant with decreased mitochondrial area consistent with its increased TBC1 recruitment. By contrast, co-expression of the 5LA variant did not show increased mitochondrial fragmentation with YFP-TBC1D15 consistent with 5LA’s defective ability to support TBC1 recruitment. These results show that perturbations to the SKY insert can be rescued by TBC1D15 expression and supports a role for TBC1D15 in FIS1-driven changes in mitochondrial morphology.

**Figure 5.**
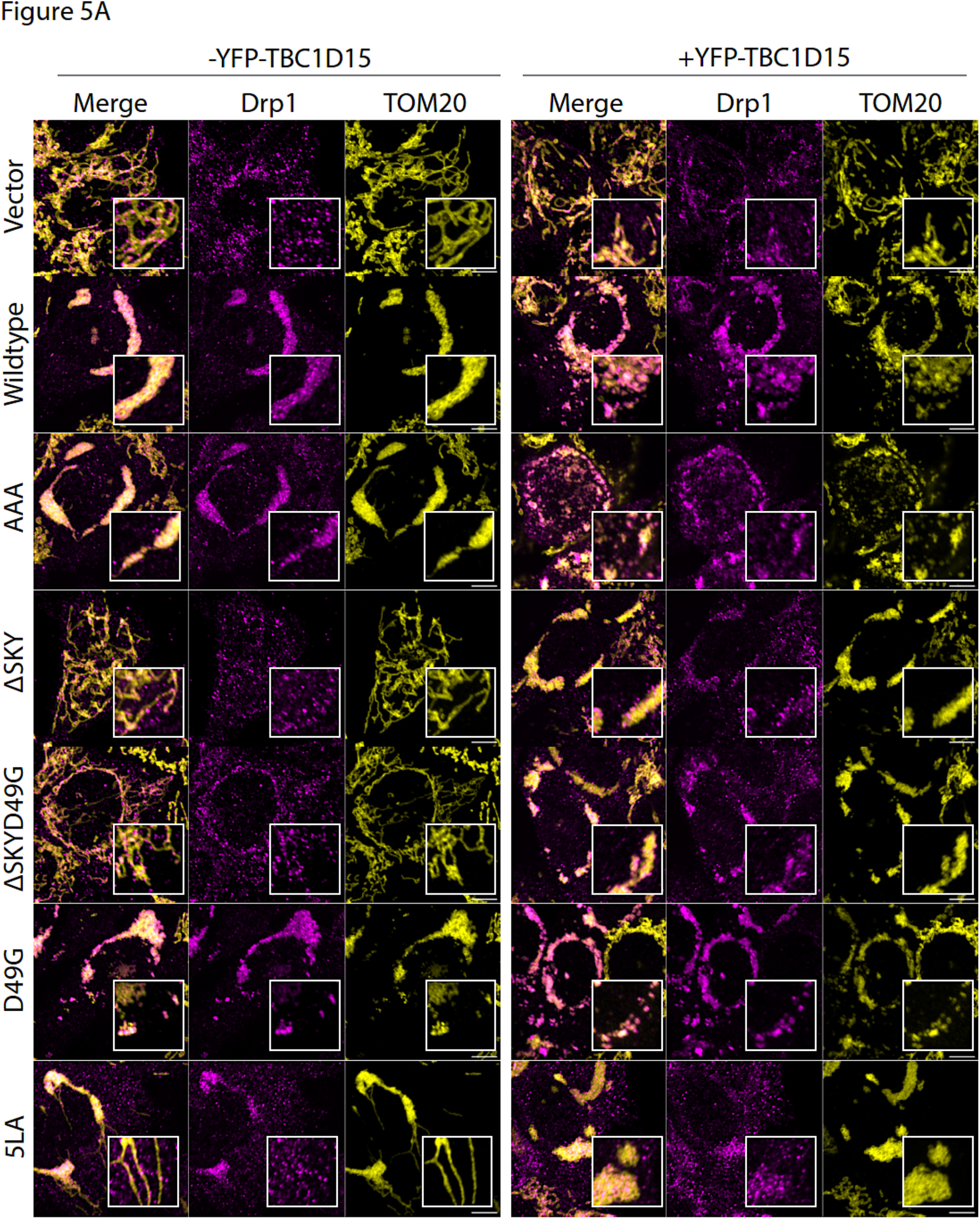
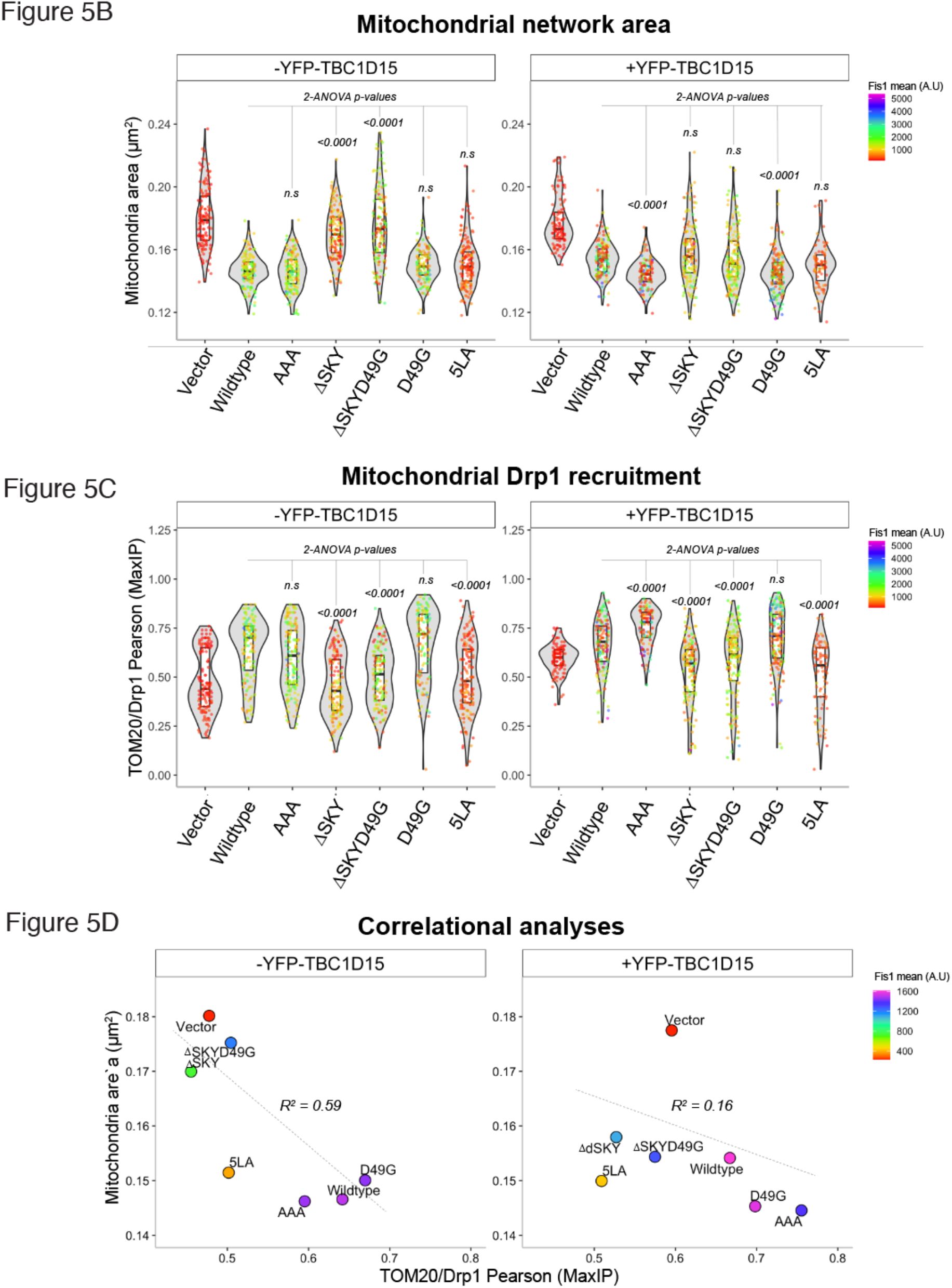
DRP1 recruitment L.O.F is partially rescued by TBC1D15 overexpression. Analyses of HCT116 cells co-overexpressing FIS1 and YFP-TBC1D15 from Figure 4 for DRP1 localization. **A,** Representative confocal images showing merged anti-DRP1 (magenta) and anti-TOM20 (yellow) from single channel images as indicated before (left panel) and after (right panel) transfection with YFP-TBC1D15. Scale bar = 100μm, insets are enlarged 4X with fluorescence intensities adjusted for clarity. **B,** Violin plots of average mitochondrial component area. **C**, Violin plot of the colocalization between mitoYFP and DRP1 from single cell maximum intensity projections was measured using Pearson’s correlation coefficient. **D,** Correlation plot to determine the relationship between mitochondrial component area and mitochondrial DRP1. Each point in **B,C** represents a single cell and each circle in **D** represents the population means and are colored based on the FIS1 expression levels determined from mean fluorescence intensity per cell. Data represent three biological replicates with *p* values calculated from 2-WAY ANOVA analyses followed by TUKEY honest significant differences (HSD).

To determine whether this rescue depended on DRP1, we immunostained for endogenous DRP1 in these experiments and determined mitochondrial co-localization (**Figures 5A, C**). Across all conditions, expression of TBC1D15 increased mitochondrial localization of DRP1. This is perhaps most notable for vector condition with endogenous FIS1 that showed a bimodal distribution of DRP1 recruitment in the violin plot with a population of cells that poorly recruited DRP1 that is abolished upon TBC1D15 expression (**Figure 5C**). In the presence of TBC1D15, the AAA variant increased DRP1 recruitment over wildtype consistent with its more pronounced effect on mitochondrial morphology with decreased mitochondrial area. D49G showed a similar effect although not statistically significant with respect to DRP1 localization. For the ΔSKY variants, we observed a bimodal distribution of DRP1 localization in the absence of TBC1D15 coexpression, which was similar to vector alone. This bimodal distribution was also eliminated upon TBC1D15 co-expression, although these ΔSKY variants still had impaired DRP1 localization compared to wildtype (**Figure 5A, C**). In fact, the extent of impaired DRP1 localization relative to wildtype decreased by ~20% regardless of TBC1D15. Similar results were found for the 5LA variant. These data indicate that expression of TBC1D15 potently rescues a fission defect in FIS1 that appears to be DRP1-dependent.

## Discussion

Here we report that mitochondrial fragmentation and impaired perinuclear clumping typical of wildtype FIS1 overexpression was abolished upon deletion of the SKY insert, which we show is a non-canonical yet highly conserved insert into the N-terminal TPR of FIS1. Both ΔSKY variants reduce DRP1 recruitment to mitochondria supporting a role for FIS1 in DRP1-mediated fission. Both ΔSKY variants reduced exogenous TBC1D15 recruitment to mitochondria and were unable to support TBC1D15 assembly into punctate structures indicating that the SKY insert also supports functionally important interactions with TBC1D15. Notably, TBC1D15 expression increases mitochondrial DRP1 localization in all conditions regardless of which FIS1 construct is co-expressed, which likely explains the partial recovery of mitochondrial morphology upon coexpression with ΔSKY variants. An important role for the SKY insert in FIS1 activity is also supported by slight gain of function activities found for AAA and D49G variants.

Previously we reported that deletion of the first eight residues of FIS1, termed the FIS1 arm, impaired DRP1 localization and mitochondrial fission (41). Here we find a similar effect in HCT116 cells upon deletion of the SKY insert, but not substitution of these residues with AAA. Both arm and SKY deletions potently impair FIS1 activity and mitochondrial DRP1 recruitment. Interestingly, we also noted that both arm and SKY deletions prevented ectopic TBC1D15 puncta formation; instead, TBC1D15 was uniformly sequestered on mitochondrial networks indicating that the FIS1 arm does not directly mediate binding, but likely regulates other interactions necessary for TBC1D15 puncta formation (41) (**Figure 4).** These observations are likely connected: Molecular dynamics simulations show intramolecular, bifurcated hydrogen-bonding between the carboxamide of Asn6 in the FIS1 arm and the backbone atoms of the SKY insert are possible (41). Such interactions would be expected to be supported by AAA and D49G, but not ΔSKY variants. NMR chemical shift changes in arm residues upon deletion of SKY also support the possibility of arm-SKY intramolecular interactions (**Figure 2**). Moreover, the NMR data for ΔSKYD49G shows conformational heterogeneity that it relieved upon deletion of the FIS1 arm (**Figure S1**) indicating that the arm is responsible for this heterogeneity; it is likely indiscriminately sampling non-native interactions with the TPR core in the absence of the SKY insert. The thermal unfolding data are also consistent with this interpretation as arm deletion restores the T_m_ to 73.6 ± 0.5 (not shown). Thus, multiple lines of evidence support that FIS1 activity requires intramolecular arm-SKY interactions that might govern the recruitment and assembly of effector proteins like TBC1D15 and DRP1 (**Figure 6**).

**Figure 6.**
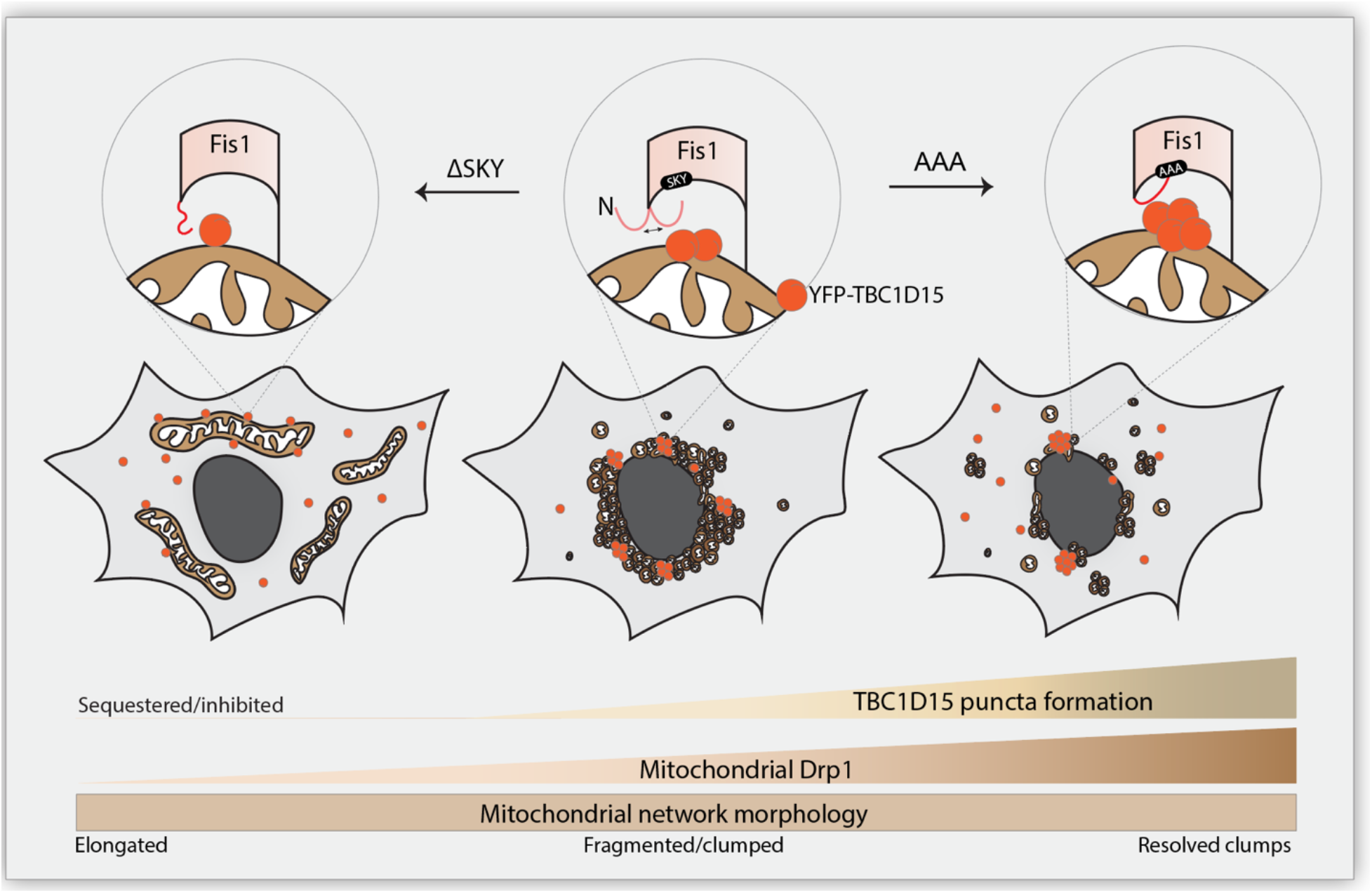
The conserved SKY insert helps to govern recruitment of DRP1 and TBC1D15 in fission. The three-residue insert in FIS1 is conserved across species for DRP1 and species-specific adaptor recruitment via conserved interactions with the FIS1 arm.

Ectopic expression of YFP-TBC1D15 increases DRP1 localization under all conditions tested including endogenous conditions, which partially rescues the mitochondrial fission defects caused by ectopic expression of FIS1 ΔSKY variants (**Figure 5**). These data indicate that TBC1D15 can drive mitochondrial fission via endogenous FIS1 and/or mechanisms that are FIS1-independent since TBC1D15 also physically interacts with DRP1(45). In this sense, these data are highly reminiscent of the Fis1p-Mdv1p-Dnm1p apparatus in yeast, where deletion of the FIS1 arm can be rescued upon Mdv1p overexpression, which also has known interactions with Dnm1p the yeast DRP1 ortholog (8). Thus, it is tempting to speculate that TBC1D15 is a functional Mdv1p ortholog in vertebrates. However, Mdv1p and TBC1D15 only share 21% sequence identity and share no discernible structural homology based on AlphaFold predictions except for disordered regions supporting the idea that FIS1 likely has species-specific adaptors.

Ectopic expression of YFP-TBC1D15 also significantly reduced mitochondrial clumps caused by FIS1 overexpression as witnessed by reduced mitochondrial area (**Figure 5**). YFP-TBC1D15 expression also appeared to stabilize FIS1 levels (**Figure S3**) thus tying TBC1D15 activity to FIS1 turnover as previously reported (30). One plausible explanation is that TBC1D15 increases mitophagic flux thus reducing clumps **(Figure 5**). Interestingly, co-expression of YFP-TBC1D15 failed to reduce the expression levels of ΔSKY variants indicating that FIS1 turnover is impaired when the SKY insert is mutated (**Figure S3**). FIS1 turnover is likely regulated by post-translational modifications, with ubiquitination playing a central role (46–49). FIS1 turnover in lipogenic cells is post-translationally regulated by stepwise acetylation and ubiquitination of unknown lysine residues that may include K46 of the SKY insert (47). Consistent with this, we note that AAA expression was significantly higher compared to when the wildtype protein was expressed (**Figure S2**).

TBC1D15 is reported to have oncogenic, lysosomal, and mitophagic functions (31, 50–54) presumably by interacting with p53-NUMB, FIS1-DRP1, and Rab7A (30, 45, 50, 51). Interestingly, FIS1’s mitophagic functions appear to be closely linked to TBC1D15-dependent tethering of Rab7A+ subcellular compartments to mitochondrial sites (31, 53). Functional links between FIS1 and TBC1D15 are demonstrated by knockout studies showing synergistic increases in mitochondrial elongation upon ablating both FIS1 and TBC1D15 (30). As such, TBC1D15 may function as a limiting factor for autophagolysosomal fusion mediated by Rab7A during mitophagy (31, 32). Interestingly, genetic ablation of FIS1 or TBC1D15 led to the formation of large LC3B structures that is indicative of impaired autophagy, providing further evidence of functional links between FIS1 and TBC1D15 (31). Our work here extends these observations by showing a new role for TBC1D15 in facilitating FIS1-mediated DRP1 recruitment. TBC1D15 is actively degraded during nutrient starvation (50), a stressor that also triggers mitochondrial elongation in mammalian cells (55), which in light of these results may derive from reduced FIS1-mediated fission.

## Experimental procedures

### Structural and phylogenetic sequence alignments

We searched PROSITE for human proteins containing TPRs (56). Putative TPR sequences alone from these proteins were then manually compiled as a FASTA formatted file and aligned on PROMALS3D using the synthetically designed TPR structure (PDB:1NA0) as a template (57). The alignment file generated by PROMALS3D was used to render the alignment figure on ESPript3.0 (58), and annotations were added to the final figure using Adobe Illustrator.

### Protein expression and purification

The soluble domains of FIS1 and variants were recombinantly expressed as SUMO protease cleavable 6xHis-smt3 fusion constructs in *E.coli BL21DE3(pRep4)* cells as previously described (59). Post-cleavage of the 6xHis-smt3 tag with recombinant SUMO, FIS1 constructs were purified to homogeneity using nickel affinity and size exclusion chromatography as described previously (59). Subsequently, samples were buffer exchanged into the final experimental buffer (100 mM HEPES pH 7.4, 200 mM NaCl, 1 mM DTT, 0.02% (w/v) sodium azide) for storage at 4 °C until biophysical analyses were conducted.

### Thermal melting assay

Thermal unfolding was monitored by light scattering and intrinsic fluorescence at 330 nm and 350 nm using a NanoTemper Prometheus instrument. Briefly, FIS1 or variants were prepared at a final concentration of 20 μM in 100 mM Hepes, pH 7.4, 200 mM NaCl, 1 mM DTT, 0.02% sodium azide. High-sensitivity capillaries (MO-K022) were then filled with each sample in four replicates for thermal scans. A melting scan was performed using an excitation power of 100%, a temperature range of 25 °C to 95 °C, and a temperature ramp of 0.5 °C/min. The resulting light scattering data were fit to a two-state model using the method of Santoro-Bolen equation (60) with the fit equation *S*(*T*)=((*S*_F_+*m*_F_\**T*)+(*S*_U_+*m*_U_\**T*)*exp(Δ*H/R*\*(1/*T*_m_-1/*T*)))/(1+exp(Δ*H/R*\*(1/*T*_m_-1/*T*))) to determine the midpoint of the unfolding transition, *T*_m_, and rendered as box and whisker plots using R.

### NMR spectroscopy

Two-dimensional ^1^H,^15^N heteronuclear single quantum coherence (HSQC) data were collected in 3 mm NMR tubes (Bruker) on a 14.1 T Bruker Avance II spectrometer equipped with a 5 mm TCI cryoprobe with a z-axis gradient. Data were collected on 100 μM ^15^N-FIS1 in 100 mM HEPES pH 7.4, 200 mM NaCl, 1 mM DTT, 0.02% (w/v) sodium azide, 10% ^2^H_2_O, 25 °C with eight transients, and 1024 (t2) × 300 (t1) complex points with acquisition times of 51.2 ms (^1^H) and 75.0 ms (^15^N). Spectra were processed with NMRPipe and analyzed with NMRAnalysis 2.5.2 (61) and NMRAssign 3.0 (62) using NMRBox (63). Chemical shift assignments for FIS1(1–125) have been previously reported (59) and for SKY variants were completed by visual inspection.

### Cell culture & Transfections

Human colorectal carcinoma cells (HCT116 cells, ATCC) were cultured in Mcoy5A supplemented with 10 mM glutamine, 10% FBS, and 1% NEAA. See table of reagents in supporting information for full details of chemicals and suppliers. Transfections were carried out in media supplemented with 2% FBS. For transfections, cells were plated on sterilized No. 1.5 glass bottom 24-well dishes (Cellvis). Optimal adherence and confluence were achieved by seeding cells at 20% confluence 48h prior to transfection. Before transfection, cell media was changed to fresh media containing 2% FBS. For transfections, plasmid DNA was added to Opti– MEM and briefly mixed by vortexing. The transfection reagent, Avalanche–Omni, was briefly vortexed and then 1μl was added to the DNA:Opti–MEM mixture (1.25ug:250μl), immediately followed by vortexing for an additional 5 sec. After 15 mins of incubation at RT, 100μl of formed transfection complexes were added dropwise into each well. Cells were incubated in transfection reagent for 6-8 hours, then changed to fresh media and incubated overnight. Cells were subsequently processed for immunofluorescence 18-24h post-transfection.

### Immunofluorescence staining

18-24h post-transfection, the medium was aspirated and replaced with 4% paraformaldehyde (prewarmed to 37°C) and incubated with gentle shaking at room temperature for 25 – 30 minutes (see table of reagents in supporting information for details). The fixative was removed and replaced with PBS. Following fixation, the cells were permeabilized by incubating with PBS/0.15% Triton X–100 for 15 min, followed by a brief wash in PBS and incubation with blocking solution (0.3% BSA/0.3% Triton X-100/PBS) for 1 hour. Cells were then incubated overnight with primary antibody mix/5% normal goat serum/blocking solution, washed three times in PBS, incubated for 1 hr with secondary antibody/blocking solution, and washed 2X in PBS/ 0.05% TWEEN 20 and once in PBS. To minimize antibody cross-reactivity in dual-labeling experiments, antibody incubations were processed sequentially, first for DRP1 (1:100) or Tom20 (1:500), followed by FIS1 (1:200).

### Image acquisition, colocalization, fluorescence intensity, and mitochondrial area analyses

Cells were visualized using a Nikon spinning-disk confocal microscope (see reagent table for detailed information). For morphology counts cells were visualized using a 100x oil objective at 0.2-micron z-slices and 0.07-micron resolution and assessed by eye for the indicated morphology. Representative confocal images were acquired and processed using FIJI. All IF-based recruitment experiments were repeated three times and at least 30 cells or more (or a total of 100 or more cells) per experimental condition were manually cropped for statistical analyses. For colocalization analysis, the FIJI coloc2 plugin was used to calculate Pearson’s Correlation between endogenous DRP1 and mitoYFP, DRP1 and TOM20, or YFP-TBC1D15 and endogenous Tom20 as described (41). A FIJI macro was used for cellular analyses and single channel/single cell z-stack images generated from MitoGraph pre-processing for the coloc2 analysis as described (41). Maximum intensity projection image stacks and images from MitoGraph preprocessing were used to measure the mean intensity of FIS1 within each cell. R was used to compile Pearson coefficients and combined in a merged data set with the MitoGraph metrics and FIS1 fluorescence intensity analysis as described (41). For analyses of YFP-TBC1D15 signal transition, YFP fluorescence intensity analyses were similarly performed in batch mode on FIJI using MitoGraph preprocessing cropped images to determine YFP mean and mode values per cell. Violin plots and ANOVA statistical calculations were also performed using R.

Batch mode pre-processing of images for mitochondrial area assessment by MitoGraph was done using R scripts previously described (41, 64). MitoGraph segmentation and noise removal were performed on cropped TIFF files using the following commands for segmentation: MitoGraph -xy 0.07 -z 0.2 -adaptive 10 -path cells. The resulting PNG files were compiled using an ImageJ macro and screened for accurate mitochondrial segmentation as previously described (41). The average mitochondrial area was then determined by multiplying the average edge length and average width values generated by MitoGraph. Mitochondrial area data was merged with mean fluorescence intensity values of FIS1 for statistical evaluation using R.

### Western Blot by JESS

Transfected HCT116 cells were harvested using a RIPA lysis kit (ProteinSimple CBS401) and cleared supernatants were saved at −20C until analyses. Capillary electrophoresis experiments were carried out using a JESS system (Protein Simple) with the 25 capillary 12-230kDa Separation module (ProteinSimple SM-W004), FIS1 antibody (Proteintech 10955-1-AP) and the Anti-Rabbit Detection Module (ProteinSimple DM-001). Setup and analysis were performed according to the manufacturer’s instructions. Briefly, samples were diluted to a final concentration of 0.2mg/ml in 0.1X sample buffer and 5X fluorescent master mix. The biotinylated ladder and the samples are then heated at 95 °C for 5 min. Once all reagents were dispensed, the plate is covered and centrifuged for 5 min at 1k rpm. Runs were performed using the instrument default settings in the Compass software (ProteinSimple, version 6.1.0). Once the run is complete, we use the Compass software to determine the signal area for each antibody. For area calculations, we use the dropped lines option. We additionally performed a total protein assay for loading level normalization using the Total Protein Detection Module (DM-TP01). The total protein area for FIS1 was normalized to overexpressed wildtype FIS1 and plotted for comparison.

## Data availability

All R scripts used for data analysis and visualization are available upon request and/or for download at https://github.com/Hill-Lab/. Raw data is available upon request.

## Supporting Information

This article contains supporting information

## Acknowledgments

Special thanks to Dr. Megan Harwig for her continued mentorship, helpful suggestions, as well as for writing all scripts and FIJI macros that were used for image-based analyses. Thanks to Florin Saitis for his assistance during sample preparation for thermal assay. Special thanks to Hill lab members, Kyle Ross and Kelsey Nolden for their useful feedback, and for proofreading the manuscript.

## Author contributions

UKI conducted all cell and IF-based experiments, analyses, and data generation. UKI and RM collected samples for WB. RT and RM conducted and analyzed JESS experiments and generated data. UKI and RBH collected and analyzed NMR and MST data. UKI and RBH conceptualized, wrote, and edited the final manuscript.

## Funding and additional information

This project was supported by the following National Institutes of Health grant R01GM067180 (to R. B. H.). The content is solely the responsibility of all authors and does not necessarily represent the official views of the federal government or the National Institutes of Health.

## Supporting Information

**FigureS1:**
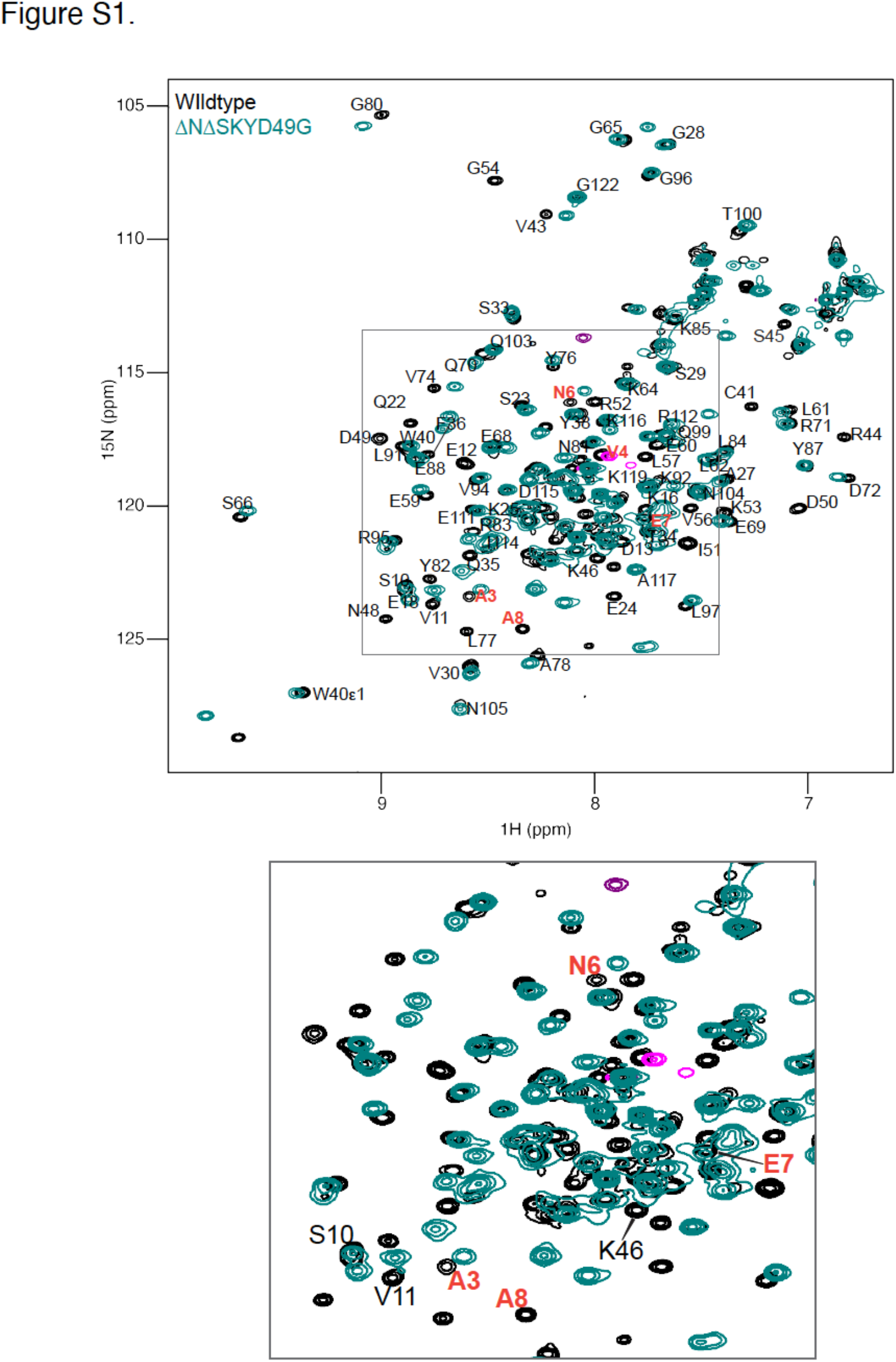
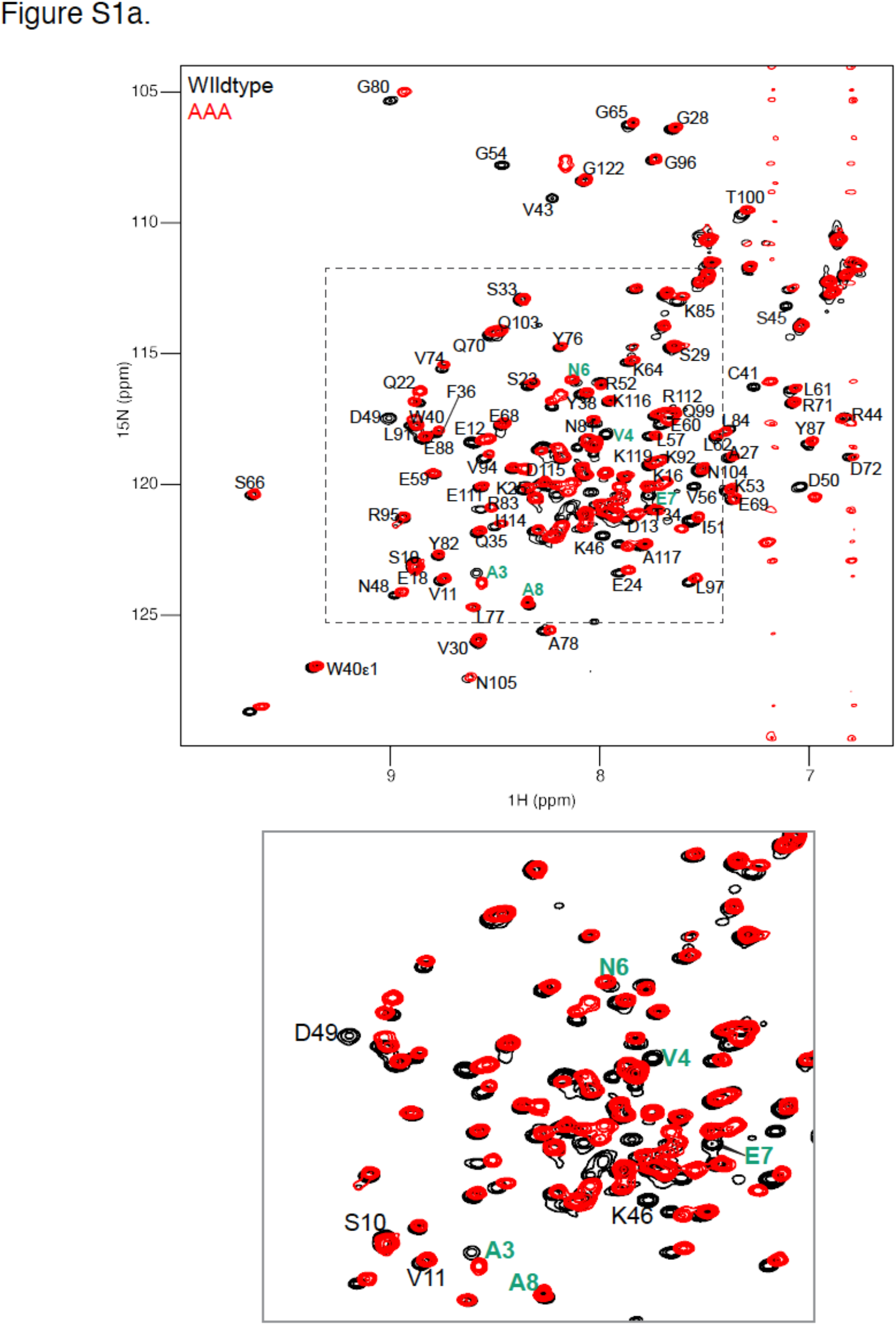
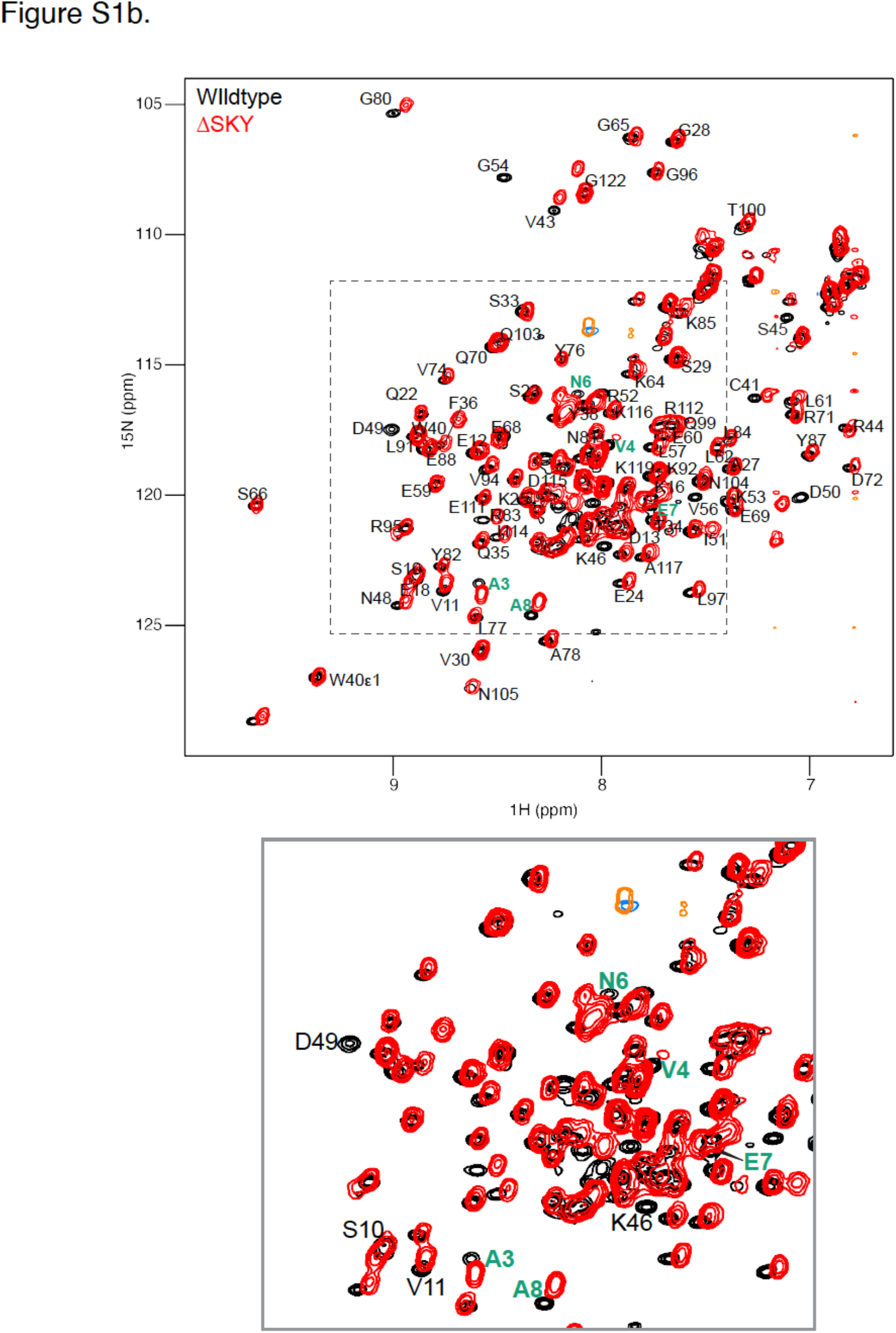
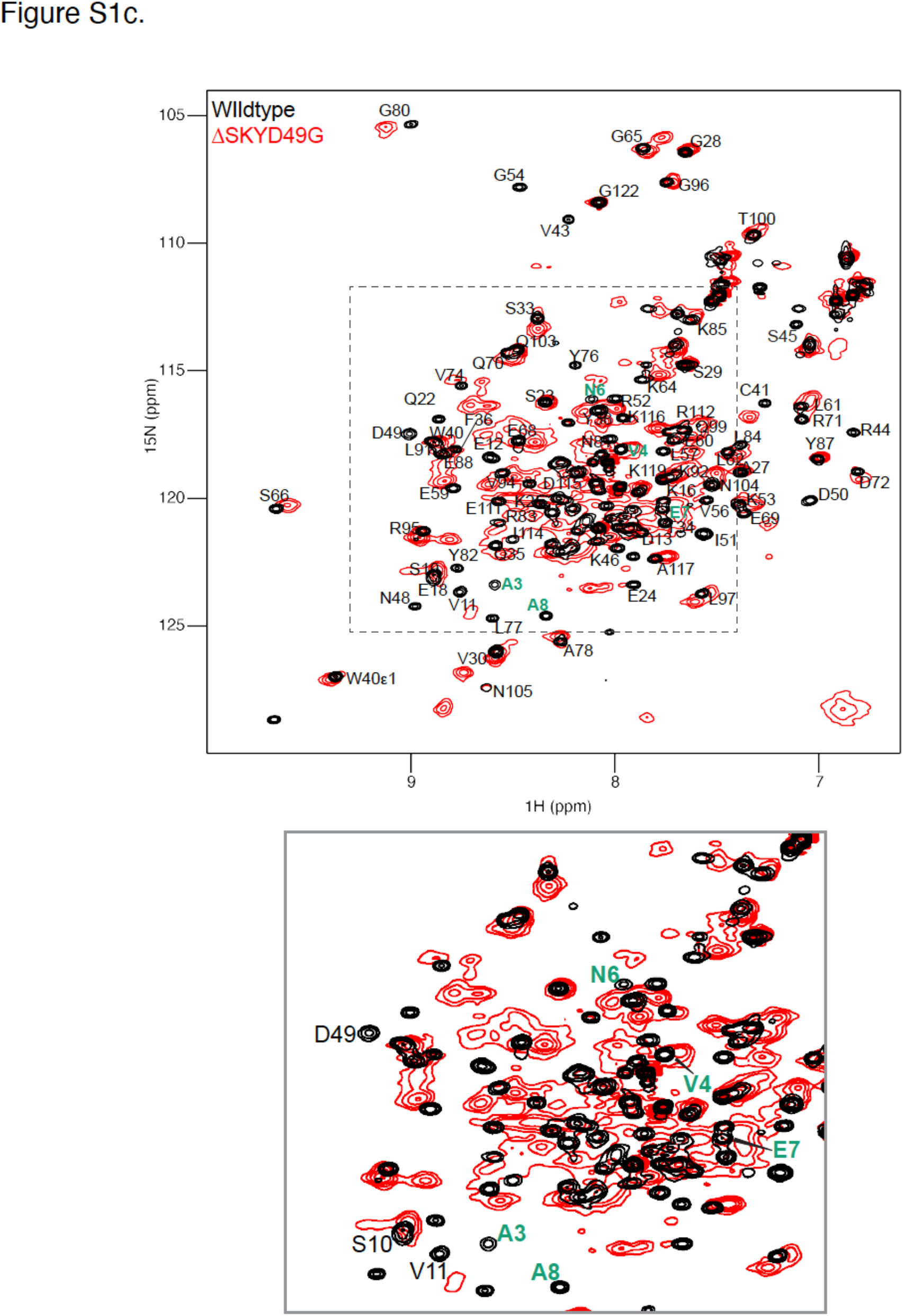
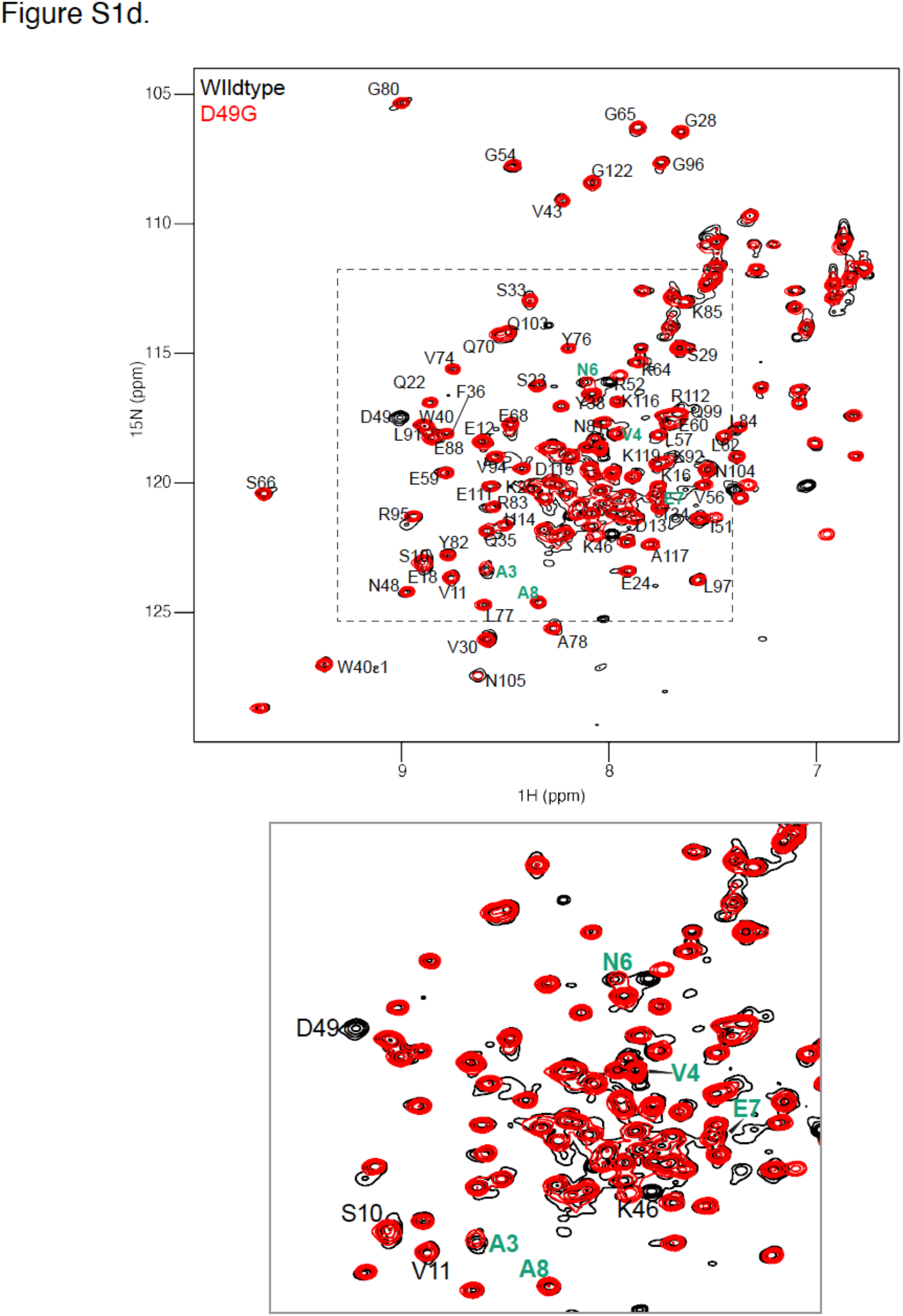
HN-HSQC spectra overlay of the cytosolic domains of wildtype (black) and ΔNΔSKYD49G (green) showing peak broadening observed in ΔSKYD49G (see **Figure S1C**) is significantly reduced upon removing amino acids (aa. 1-8) that make up the FIS1 “arm” region. Arm residue peaks are highlighted (in red) in enlarged region shown below the full spectra. **Figure S1a–S1d:** Shows HN-HSQC spectra overlays for wildtype FIS1 (black) against SKY variants (in red), AAA, ΔSKY, ΔSKYD49G, and D49G respectively. Arm residue peaks are highlighted (in green) in enlarged region shown below the full spectra.

**Figure S2:**
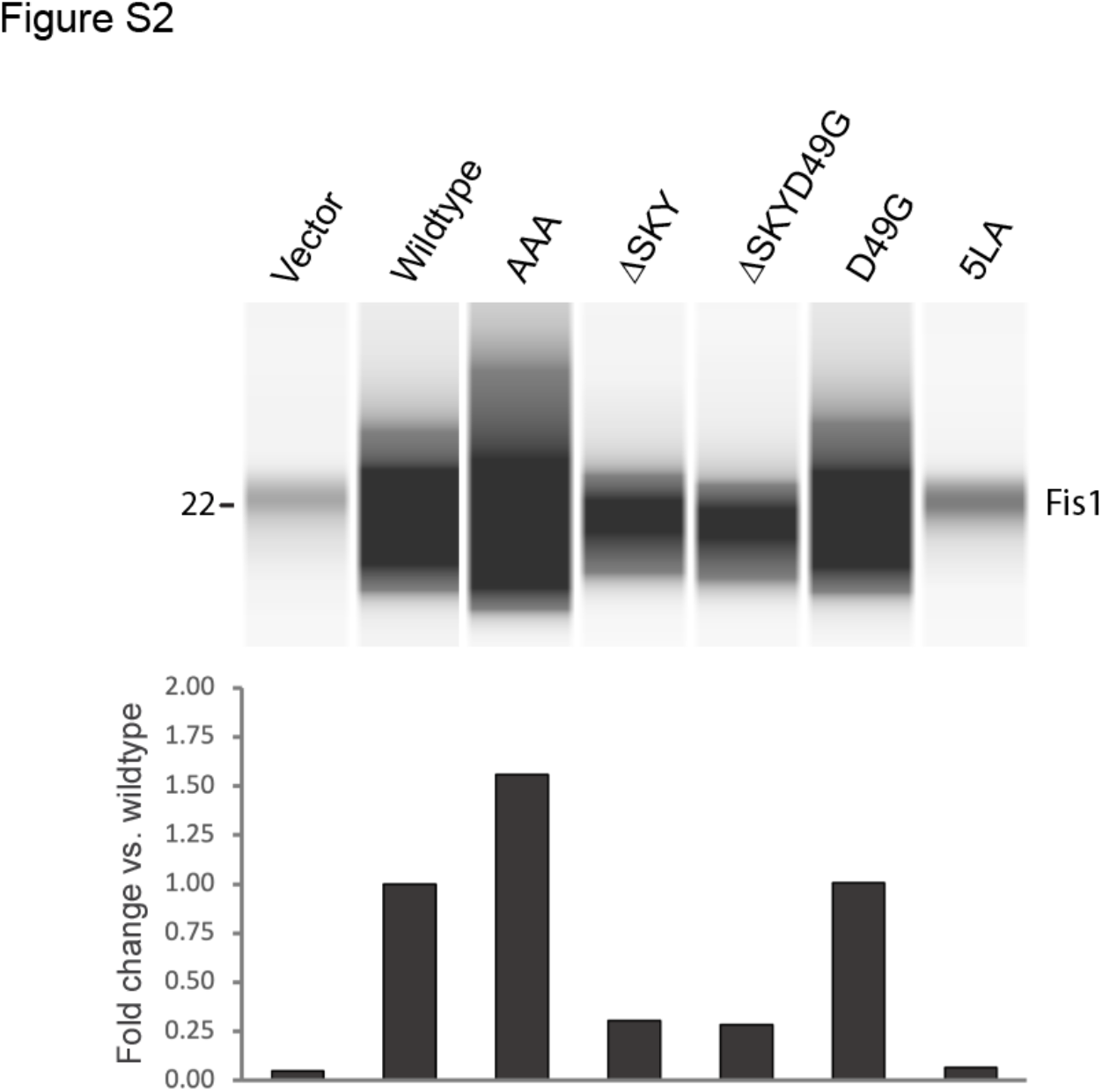
Steady-state expression of exogenously expressed FIS1 was analyzed by Western blot of whole cell lysates harvested from HCT116 cells co-transfected with mitoYFP and vector/FIS1. The total signal area for FIS1 was then quantified as normalized values compared to exogenous wildtype FIS1 expression. See the experimental procedure section for a comprehensive outline of method.

**Figure S3:**
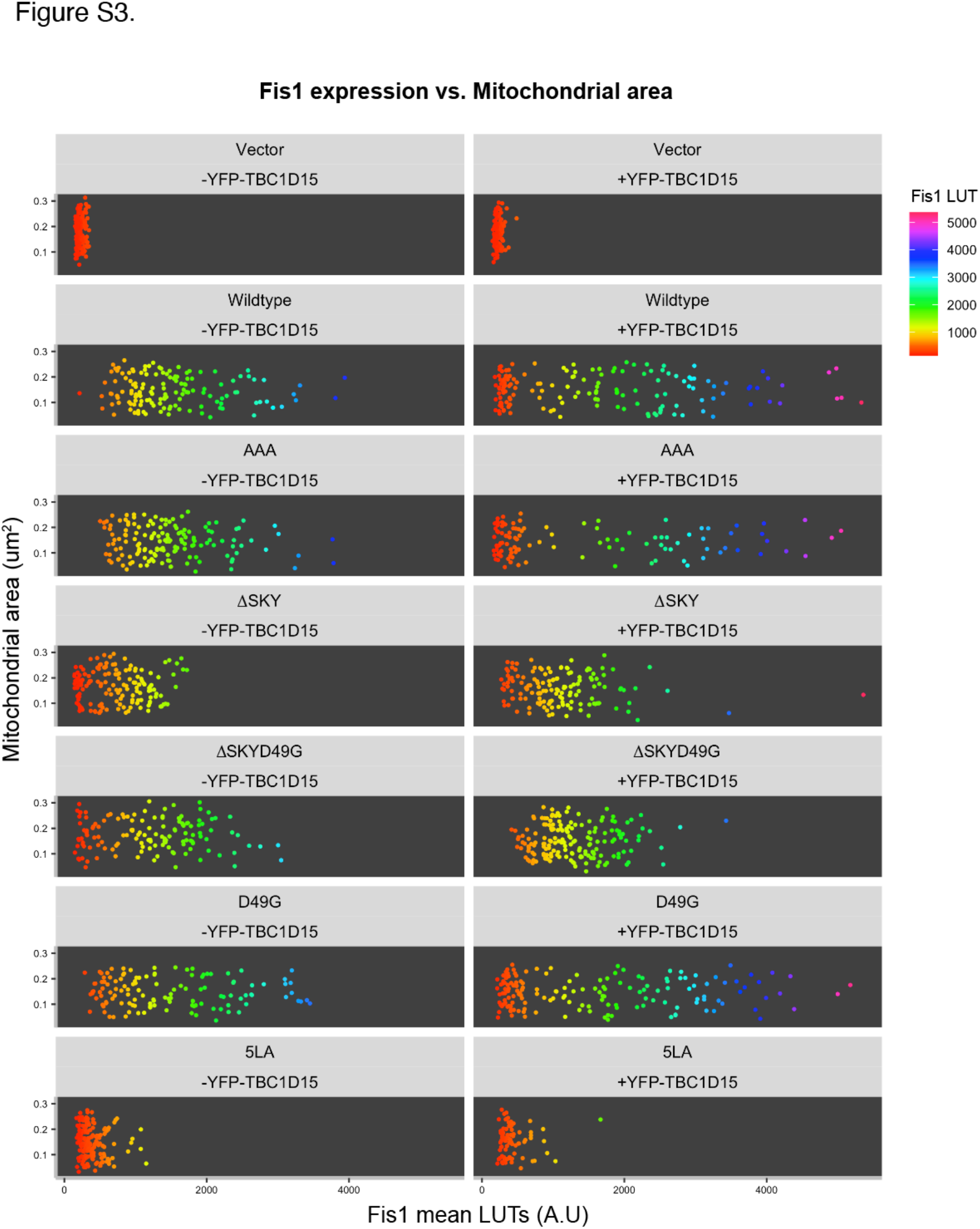
HCT116 cells were co-transfected without (-) or with (+) YFP-TBC1D15 and wildtype FIS1 or SKY variants. Mitochondrial network area (y-axis) and FIS1 mean fluorescence intensities (x-axis) were measured and plotted for each cell.

## Table of Reagents

List of reagents used in this study. Source and additional notes are included as needed.

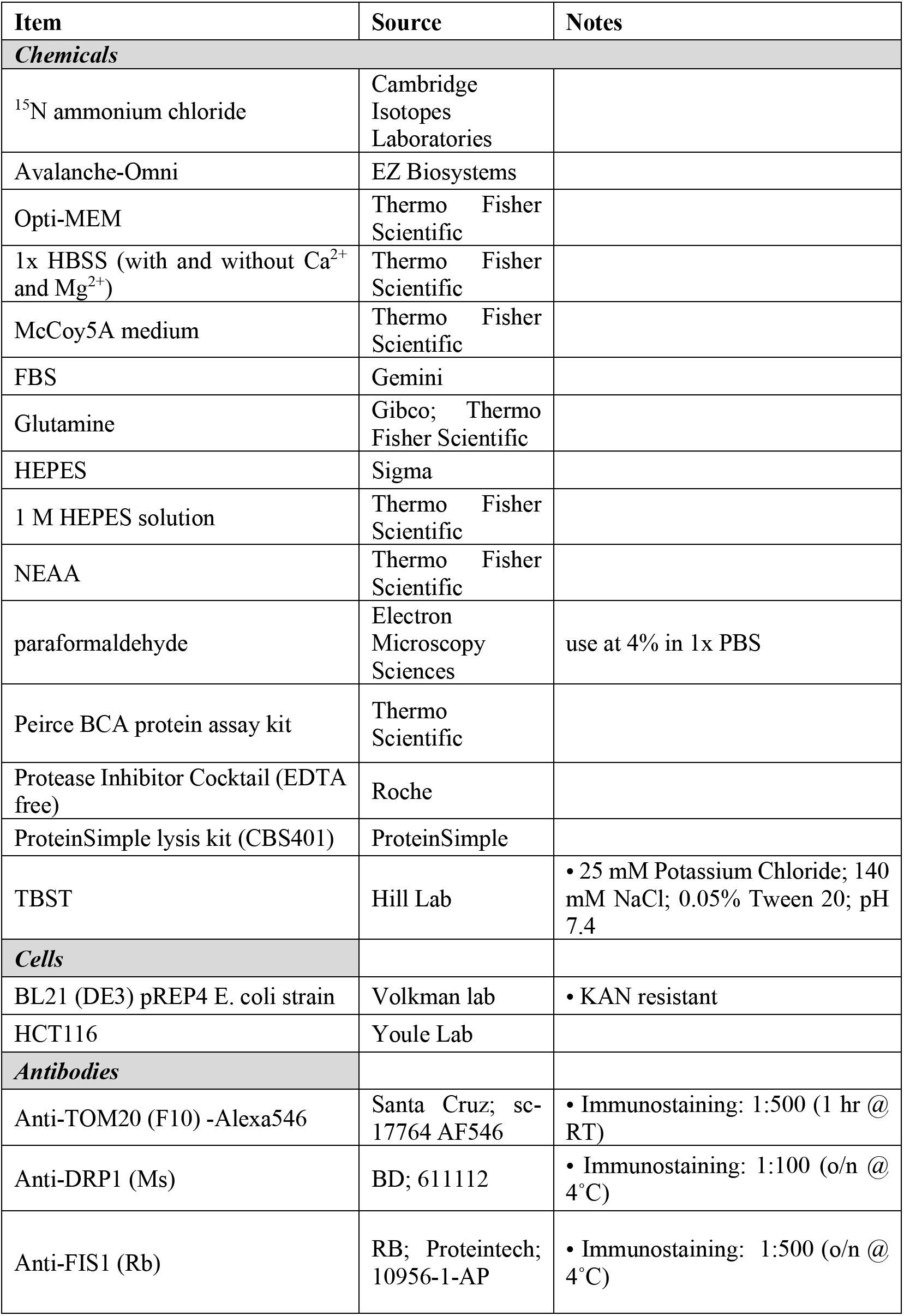
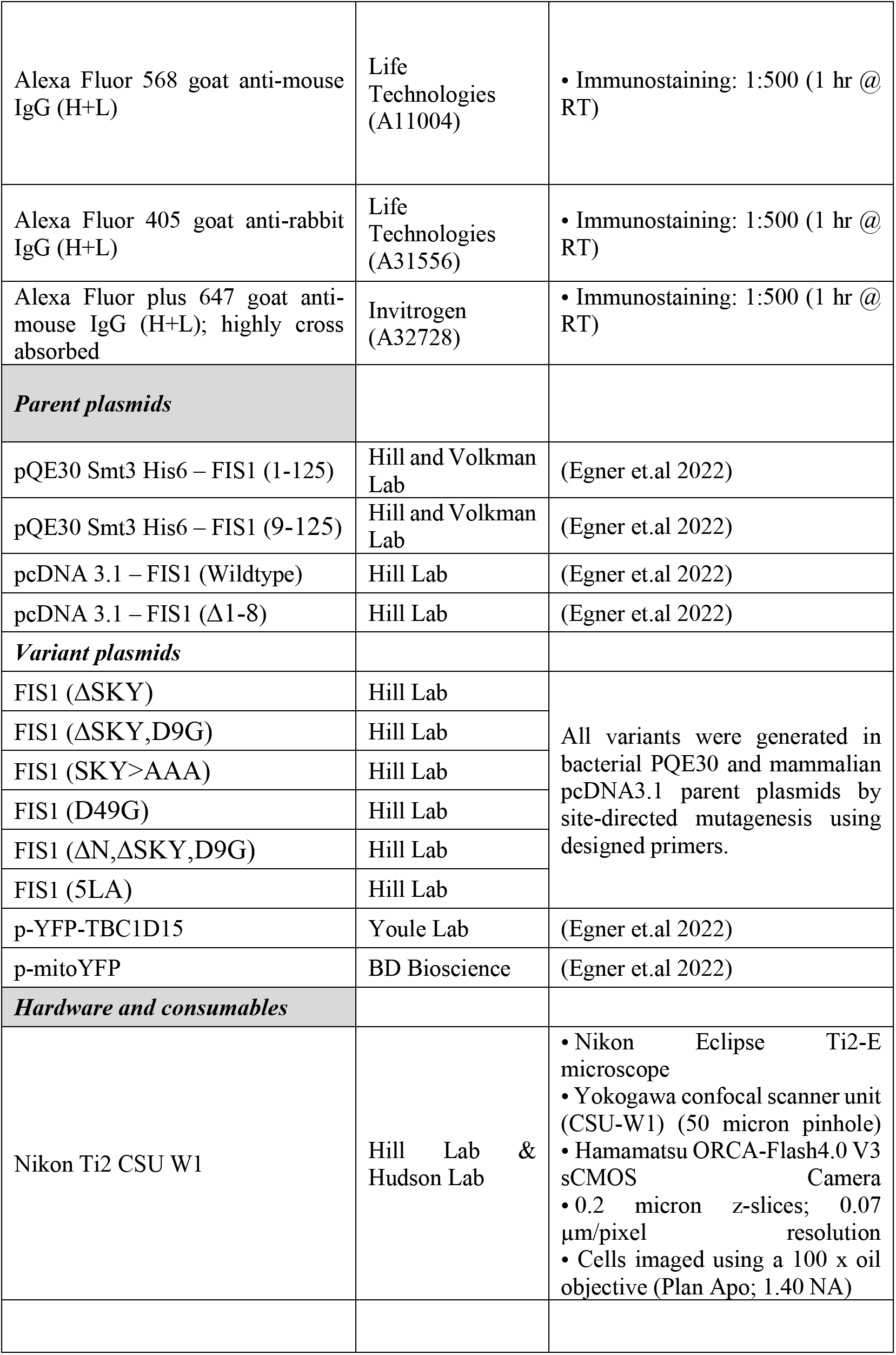
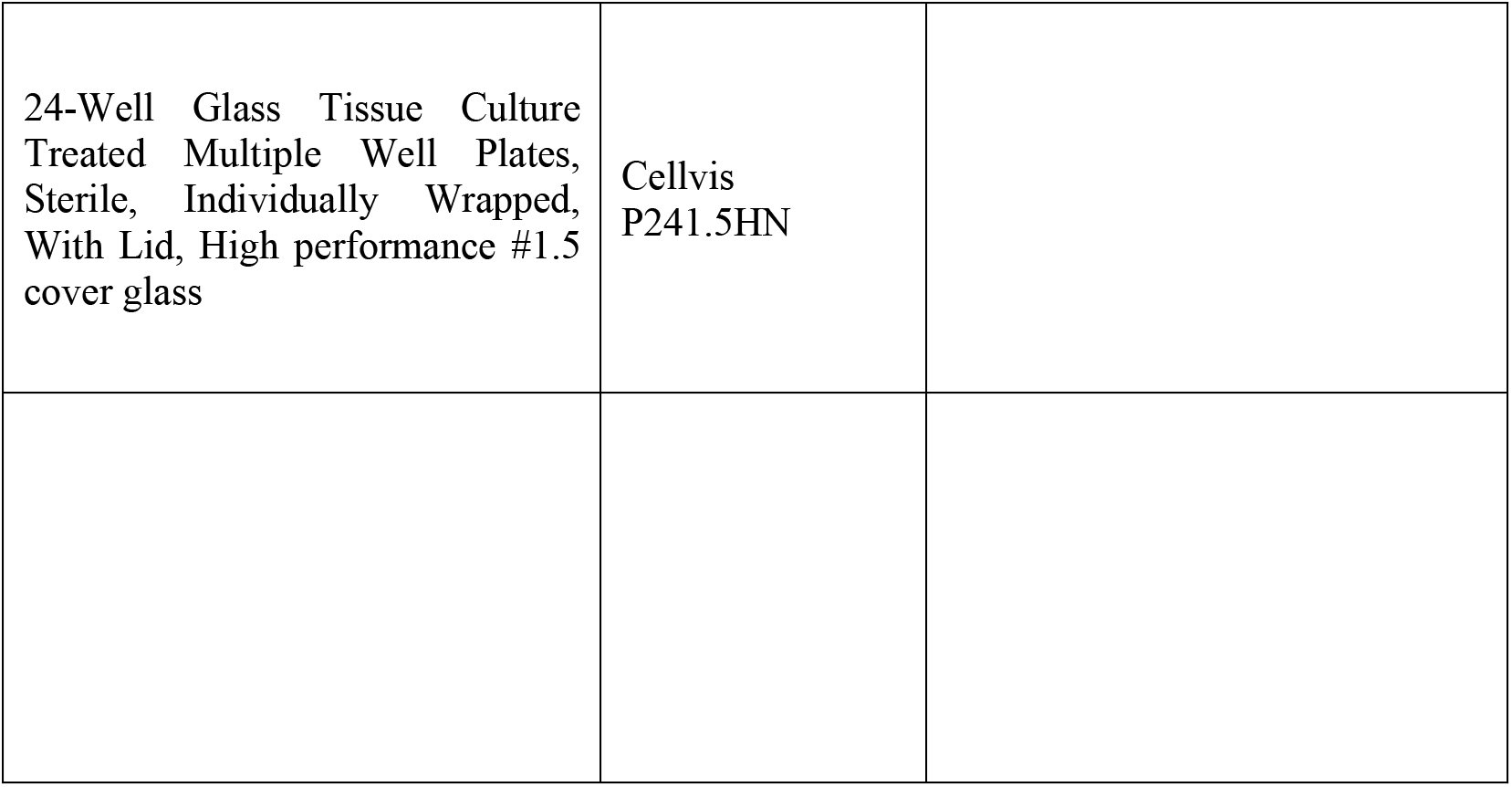

| Item | Source | Notes |
| --- | --- | --- |
| **Chemicals** |  |  |
| <sup>15</sup>N ammonium chloride | Cambridge Isotopes Laboratories |  |
| Avalanche-Omni | EZ Biosystems |  |
| Opti-MEM | Thermo Fisher Scientific |  |
| 1x HBSS (with and without Ca<sup>2+</sup> and Mg<sup>2+</sup>) | Thermo Fisher Scientific |  |
| McCoy5A medium | Thermo Fisher Scientific |  |
| FBS | Gemini |  |
| Glutamine | Gibco; Thermo Fisher Scientific |  |
| HEPES | Sigma |  |
| 1 M HEPES solution | Thermo Fisher Scientific |  |
| NEAA | Thermo Fisher Scientific |  |
| paraformaldehyde | Electron Microscopy Sciences | use at 4% in 1x PBS |
| Peirce BCA protein assay kit | Thermo Scientific |  |
| Protease Inhibitor Cocktail (EDTA free) | Roche |  |
| ProteinSimple lysis kit (CBS401) | ProteinSimple |  |
| TBST | Hill Lab | • 25 mM Potassium Chloride; 140 mM NaCl; 0.05% Tween 20; pH 7.4 |
| **Cells** |  |  |
| BL21 (DE3) pREP4 E. coli strain | Volkman lab | • KAN resistant |
| HCT116 | Youle Lab |  |
| **Antibodies** |  |  |
| Anti-TOM20 (F10) -Alexa546 | Santa Cruz; sc-17764 AF546 | • Immunostaining: 1:500 (1 hr @ RT) |
| Anti-DRP1 (Ms) | BD; 611112 | • Immunostaining: 1:100 (o/n @ 4°C) |
| Anti-FIS1 (Rb) | RB; Proteintech; 10956-1-AP | • Immunostaining: 1:500 (o/n @ 4°C) |
| Alexa Fluor 568 goat anti-mouse IgG (H+L) | Life Technologies (A11004) | • Immunostaining: 1:500 (1 hr @ RT) |
| Alexa Fluor 405 goat anti-rabbit IgG (H+L) | Life Technologies (A31556) | • Immunostaining: 1:500 (1 hr @ RT) |
| Alexa Fluor plus 647 goat anti-mouse IgG (H+L); highly cross absorbed | Invitrogen (A32728) | • Immunostaining: 1:500 (1 hr @ RT) |
| **Parent plasmids** |  |  |
| pQE30 Smt3 His6 – FIS1 (1-125) | Hill and Volkman Lab | (Egner et.al 2022) |
| pQE30 Smt3 His6 – FIS1 (9-125) | Hill and Volkman Lab | (Egner et.al 2022) |
| pcDNA 3.1 – FIS1 (Wildtype) | Hill Lab | (Egner et.al 2022) |
| pcDNA 3.1 – FIS1 (Δ1-8) | Hill Lab | (Egner et.al 2022) |
| **Variant plasmids** |  |  |
| FIS1 (ΔSKY) | Hill Lab | All variants were generated in bacterial PQE30 and mammalian pcDNA3.1 parent plasmids by site-directed mutagenesis using designed primers. |
| FIS1 (ΔSKY,D9G) | Hill Lab | |
| FIS1 (SKY>AAA) | Hill Lab |  |
| FIS1 (D49G) | Hill Lab |  |
| FIS1 (ΔN,ΔSKY,D9G) | Hill Lab | |
| FIS1 (5LA) | Hill Lab |  |
| p-YFP-TBC1D15 | Youle Lab | (Egner et.al 2022) |
| p-mitoYFP | BD Bioscience | (Egner et.al 2022) |
| **Hardware and consumables** |  |  |
| Nikon Ti2 CSU W1 | Hill Lab & Hudson Lab | • Nikon Eclipse Ti2-E microscope<br>• Yokogawa confocal scanner unit (CSU-W1) (50 micron pinhole)<br>• Hamamatsu ORCA-Flash4.0 V3 sCMOS Camera<br>• 0.2 micron z-slices; 0.07 μm/pixel resolution<br>• Cells imaged using a 100 x oil objective (Plan Apo; 1.40 NA) |
| 24-Well Glass Tissue Culture Treated Multiple Well Plates, Sterile, Individually Wrapped, With Lid, High performance #1.5 cover glass | Cellvis P241.5HN |  |

## References

1. Liesa, M., and Shirihai, O. S. (2013) Mitochondrial dynamics in the regulation of nutrient utilization and energy expenditure. Cell Metab. 17, 491–506

2. Galloway, C. A., Lee, H., and Yoon, Y. (2012) Mitochondrial morphology-emerging role in bioenergetics. Free Radic. Biol. Med. 53, 2218–2228

3. Mishra, P., and Chan, D. C. (2016) Metabolic regulation of mitochondrial dynamics. J. Cell Biol. 212, 379–387

4. Picard, M., and Shirihai, O. S. (2022) Mitochondrial signal transduction. Cell Metab. 34, 1620–1653

5. Sesaki, H., and Jensen, R. E. (1999) Division versus fusion: Dnm1p and Fzo1p antagonistically regulate mitochondrial shape. J. Cell Biol. 147, 699–706

6. Jakobs, S., Martini, N., Schauss, A. C., Egner, A., Westermann, B., and Hell, S. W. (2003) Spatial and temporal dynamics of budding yeast mitochondria lacking the division component Fis1p. J. Cell Sci. 116, 2005–2014

7. Cerveny, K. L., and Jensen, R. E. (2003) The WD-repeats of Net2p interact with Dnm1p and Fis1p to regulate division of mitochondria. Mol. Biol. Cell. 14, 4126–4139

8. Mozdy, A. D., McCaffery, J. M., and Shaw, J. M. (2000) Dnm1p GTPase-mediated mitochondrial fission is a multi-step process requiring the novel integral membrane component Fis1p. J. Cell Biol. 151, 367–380

9. Tieu, Q., and Nunnari, J. (2000) Mdv1p is a WD repeat protein that interacts with the dynamin-related GTPase, Dnm1p, to trigger mitochondrial division. J. Cell Biol. 151, 353–366

10. Fekkes, P., Shepard, K. A., and Yaffe, M. P. (2000) Gag3p, an outer membrane protein required for fission of mitochondrial tubules. J. Cell Biol. 151, 333–340

11. Bhar, D., Karren, M. A., Babst, M., and Shaw, J. M. (2006) Dimeric Dnm1-G385D interacts with Mdv1 on mitochondria and can be stimulated to assemble into fission complexes containing Mdv1 and Fis1. J. Biol. Chem. 281, 17312–17320

12. Schauss, A. C., Bewersdorf, J., and Jakobs, S. (2006) Fis1p and Caf4p, but not Mdv1p, determine the polar localization of Dnm1p clusters on the mitochondrial surface. J. Cell Sci. 119, 3098–3106

13. Zhang, Y., and Chan, D. C. (2007) Structural basis for recruitment of mitochondrial fission complexes by Fis1. Proc. Natl. Acad. Sci. USA. 104, 18526–18530

14. Koppenol-Raab, M., Harwig, M. C., Posey, A. E., Egner, J. M., MacKenzie, K. R., and Hill, R. B. (2016) A Targeted Mutation Identified through pKa Measurements Indicates a Postrecruitment Role for Fis1 in Yeast Mitochondrial Fission. J. Biol. Chem. 291, 20329–20344

15. Yoon, Y., Krueger, E. W., Oswald, B. J., and McNiven, M. A. (2003) The mitochondrial protein hFis1 regulates mitochondrial fission in mammalian cells through an interaction with the dynamin-like protein DLP1. Mol. Cell. Biol. 23, 5409–5420

16. James, D. I., Parone, P. A., Mattenberger, Y., and Martinou, J.-C. (2003) hFis1, a novel component of the mammalian mitochondrial fission machinery. J. Biol. Chem. 278, 36373–36379

17. Jofuku, A., Ishihara, N., and Mihara, K. (2005) Analysis of functional domains of rat mitochondrial Fis1, the mitochondrial fission-stimulating protein. Biochem. Biophys. Res. Commun. 333, 650–659

18. Koch, A., Yoon, Y., Bonekamp, N. A., McNiven, M. A., and Schrader, M. (2005) A role for Fis1 in both mitochondrial and peroxisomal fission in mammalian cells. Mol. Biol. Cell. 16, 5077–5086

19. Kobayashi, S., Tanaka, A., and Fujiki, Y. (2007) Fis1, DLP1, and Pex11p coordinately regulate peroxisome morphogenesis. Exp. Cell Res. 313, 1675–1686

20. Zhang, X., and Hu, J. (2009) Two small protein families, DYNAMIN-RELATED PROTEIN3 and FISSION1, are required for peroxisome fission in Arabidopsis. Plant J. 57, 146–159

21. Scott, I., Tobin, A. K., and Logan, D. C. (2006) BIGYIN, an orthologue of human and yeast FIS1 genes functions in the control of mitochondrial size and number in Arabidopsis thaliana. J. Exp. Bot. 57, 1275–1280

22. Zhang, X., and Hu, J. (2010) The Arabidopsis chloroplast division protein DYNAMIN-RELATED PROTEIN5B also mediates peroxisome division. Plant Cell. 22, 431–442

23. Wells, R. C., Picton, L. K., Williams, S. C. P., Tan, F. J., and Hill, R. B. (2007) Direct binding of the dynamin-like GTPase, Dnm1, to mitochondrial dynamics protein Fis1 is negatively regulated by the Fis1 N-terminal arm. J. Biol. Chem. 282, 33769–33775

24. Fröhlich, C., Grabiger, S., Schwefel, D., Faelber, K., Rosenbaum, E., Mears, J., Rocks, O., and Daumke, O. (2013) Structural insights into oligomerization and mitochondrial remodelling of dynamin 1-like protein. EMBO J. 32, 1280–1292

25. Otera, H., Wang, C., Cleland, M. M., Setoguchi, K., Yokota, S., Youle, R. J., and Mihara, K. (2010) Mff is an essential factor for mitochondrial recruitment of Drp1 during mitochondrial fission in mammalian cells. J. Cell Biol. 191, 1141–1158

26. Palmer, C. S., Elgass, K. D., Parton, R. G., Osellame, L. D., Stojanovski, D., and Ryan, M. T. (2013) Adaptor proteins MiD49 and MiD51 can act independently of Mff and Fis1 in Drp1 recruitment and are specific for mitochondrial fission. J. Biol. Chem. 288, 27584–27593

27. Losón, O. C., Song, Z., Chen, H., and Chan, D. C. (2013) Fis1, Mff, MiD49, and MiD51 mediate Drp1 recruitment in mitochondrial fission. Mol. Biol. Cell. 24, 659–667

28. Osellame, L. D., Singh, A. P., Stroud, D. A., Palmer, C. S., Stojanovski, D., Ramachandran, R., and Ryan, M. T. (2016) Cooperative and independent roles of the Drp1 adaptors Mff, MiD49 and MiD51 in mitochondrial fission. J. Cell Sci. 129, 2170–2181

29. Gandre-Babbe, S., and van der Bliek, A. M. (2008) The novel tail-anchored membrane protein Mff controls mitochondrial and peroxisomal fission in mammalian cells. Mol. Biol. Cell. 19, 2402–2412

30. Onoue, K., Jofuku, A., Ban-Ishihara, R., Ishihara, T., Maeda, M., Koshiba, T., Itoh, T., Fukuda, M., Otera, H., Oka, T., Takano, H., Mizushima, N., Mihara, K., and Ishihara, N. (2013) Fis1 acts as a mitochondrial recruitment factor for TBC1D15 that is involved in regulation of mitochondrial morphology. J. Cell Sci. 126, 176–185

31. Yamano, K., Fogel, A. I., Wang, C., van der Bliek, A. M., and Youle, R. J. (2014) Mitochondrial Rab GAPs govern autophagosome biogenesis during mitophagy. Elife. 3, e01612

32. Shen, Q., Yamano, K., Head, B. P., Kawajiri, S., Cheung, J. T. M., Wang, C., Cho, J.-H., Hattori, N., Youle, R. J., and van der Bliek, A. M. (2014) Mutations in Fis1 disrupt orderly disposal of defective mitochondria. Mol. Biol. Cell. 25, 145–159

33. Ihenacho, U. K., Meacham, K. A., Harwig, M. C., Widlansky, M. E., and Hill, R. B. (2021) Mitochondrial fission protein 1: emerging roles in organellar form and function in health and disease. Front. Endocrinol. (Lausanne). 12, 660095

34. Kleele, T., Rey, T., Winter, J., Zaganelli, S., Mahecic, D., Perreten Lambert, H., Ruberto, F. P., Nemir, M., Wai, T., Pedrazzini, T., and Manley, S. (2021) Distinct fission signatures predict mitochondrial degradation or biogenesis. Nature. 593, 435–439

35. Suzuki, M., Jeong, S. Y., Karbowski, M., Youle, R. J., and Tjandra, N. (2003) The solution structure of human mitochondria fission protein Fis1 reveals a novel TPR-like helix bundle. J. Mol. Biol. 334, 445–458

36. Dohm, J. A., Lee, S. J., Hardwick, J. M., Hill, R. B., and Gittis, A. G. (2004) Cytosolic domain of the human mitochondrial fission protein Fis1 adopts a TPR fold. Proteins Struct Funct Bioinform. 54, 153–156

37. Blatch, G. L., and Lässle, M. (1999) The tetratricopeptide repeat: a structural motif mediating protein-protein interactions. Bioessays. 21, 932–939

38. Allan, R. K., and Ratajczak, T. (2011) Versatile TPR domains accommodate different modes of target protein recognition and function. Cell Stress Chaperones. 16, 353–367

39. Serasinghe, M. N., and Yoon, Y. (2008) The mitochondrial outer membrane protein hFis1 regulates mitochondrial morphology and fission through self-interaction. Exp. Cell Res. 314, 3494–3507

40. Lees, J. P. B., Manlandro, C. M., Picton, L. K., Tan, A. Z. E., Casares, S., Flanagan, J. M., Fleming, K. G., and Hill, R. B. (2012) A designed point mutant in Fis1 disrupts dimerization and mitochondrial fission. J. Mol. Biol. 423, 143–158

41. Egner, J. M., Nolden, K. A., Harwig, M. C., Bonate, R. P., De Anda, J., Tessmer, M. H., Noey, E. L., Ihenacho, U. K., Liu, Z., Peterson, F. C., Wong, G. C. L., Widlansky, M. E., and Hill, R. B. (2022) Structural studies of human fission protein FIS1 reveal a dynamic region important for GTPase DRP1 recruitment and mitochondrial fission. J. Biol. Chem.

42. Suzuki, M., Neutzner, A., Tjandra, N., and Youle, R. J. (2005) Novel structure of the N terminus in yeast Fis1 correlates with a specialized function in mitochondrial fission. J. Biol. Chem. 280, 21444–21452

43. UniProt Consortium (2023) Uniprot: the universal protein knowledgebase in 2023. Nucleic Acids Res. 51, D523–D531

44. Efimov, A. V. (1991) Structure of alpha-alpha-hairpins with short connections. Protein Eng. 4, 245–250

45. Sun, S., Yu, W., Xu, H., Li, C., Zou, R., Wu, N. N., Wang, L., Ge, J., Ren, J., and Zhang, Y. (2022) TBC1D15-Drp1 interaction-mediated mitochondrial homeostasis confers cardioprotection against myocardial ischemia/reperfusion injury. Metab. Clin. Exp. 134, 155239

46. Zhang, Q., Wu, J., Wu, R., Ma, J., Du, G., Jiao, R., Tian, Y., Zheng, Z., and Yuan, Z. (2012) DJ-1 promotes the proteasomal degradation of Fis1: implications of DJ-1 in neuronal protection. Biochem. J. 447, 261–269

47. Wang, L., Zhang, T., Wang, L., Cai, Y., Zhong, X., He, X., Hu, L., Tian, S., Wu, M., Hui, L., Zhang, H., and Gao, P. (2017) Fatty acid synthesis is critical for stem cell pluripotency via promoting mitochondrial fission. EMBO J. 36, 1330–1347

48. Waters, E., Wilkinson, K. A., Harding, A. L., Carmichael, R. E., Robinson, D., Colley, H. E., and Guo, C. (2022) The SUMO protease SENP3 regulates mitochondrial autophagy mediated by Fis1. EMBO Rep. 23, e48754

49. Yu, Y., Peng, X.-D., Qian, X.-J., Zhang, K.-M., Huang, X., Chen, Y.-H., Li, Y.-T., Feng, G.-K., Zhang, H.-L., Xu, X.-L., Li, S., Li, X., Mai, J., Li, Z.-L., Huang, Y., Yang, D., Zhou, L.-H., Zhong, Z.-Y., Li, J.-D., Deng, R., and Zhu, X.-F. (2021) Fis1 phosphorylation by Met promotes mitochondrial fission and hepatocellular carcinoma metastasis. Signal Transduct. Target. Ther. 6, 401

50. Feldman, D. E., Chen, C., Punj, V., and Machida, K. (2013) The TBC1D15 oncoprotein controls stem cell self-renewal through destabilization of the Numb-p53 complex. PLoS One. 8, e57312

51. Jongsma, M. L., Bakker, J., Cabukusta, B., Liv, N., van Elsland, D., Fermie, J., Akkermans, J. L., Kuijl, C., van der Zanden, S. Y., Janssen, L., Hoogzaad, D., van der Kant, R., Wijdeven, R. H., Klumperman, J., Berlin, I., and Neefjes, J. (2020) SKIP-HOPS recruits TBC1D15 for a Rab7-to-Arl8b identity switch to control late endosome transport. EMBO J. 39, e102301

52. Meneses-Salas, E., García-Melero, A., Kanerva, K., Blanco-Muñoz, P., Morales-Paytuvi, F., Bonjoch, J., Casas, J., Egert, A., Beevi, S. S., Jose, J., Llorente-Cortés, V., Rye, K.-A., Heeren, J., Lu, A., Pol, A., Tebar, F., Ikonen, E., Grewal, T., Enrich, C., and Rentero, C. (2020) Annexin A6 modulates TBC1D15/Rab7/StARD3 axis to control endosomal cholesterol export in NPC1 cells. Cell Mol. Life Sci. 77, 2839–2857

53. Wong, Y. C., Ysselstein, D., and Krainc, D. (2018) Mitochondria-lysosome contacts regulate mitochondrial fission via RAB7 GTP hydrolysis. Nature. 554, 382–386

54. Yu, W., Sun, S., Xu, H., Li, C., Ren, J., and Zhang, Y. (2020) TBC1D15/RAB7-regulated mitochondria-lysosome interaction confers cardioprotection against acute myocardial infarction-induced cardiac injury. Theranostics. 10, 11244–11263

55. Rambold, A. S., Kostelecky, B., Elia, N., and Lippincott-Schwartz, J. (2011) Tubular network formation protects mitochondria from autophagosomal degradation during nutrient starvation. Proc. Natl. Acad. Sci. USA. 108, 10190–10195

56. Sigrist, C. J. A., de Castro, E., Cerutti, L., Cuche, B. A., Hulo, N., Bridge, A., Bougueleret, L., and Xenarios, I. (2013) New and continuing developments at PROSITE. Nucleic Acids Res. 41, D344–7

57. Pei, J., Kim, B.-H., and Grishin, N. V. (2008) PROMALS3D: a tool for multiple protein sequence and structure alignments. Nucleic Acids Res. 36, 2295–2300

58. Robert, X., and Gouet, P. (2014) Deciphering key features in protein structures with the new ENDscript server. Nucleic Acids Res. 42, W320–W324

59. Egner, J. M., Jensen, D. R., Olp, M. D., Kennedy, N. W., Volkman, B. F., Peterson, F. C., Smith, B. C., and Hill, R. B. (2018) Development and Validation of 2D Difference Intensity Analysis for Chemical Library Screening by Protein-Detected NMR Spectroscopy. Chembiochem. 19, 448–458

60. Santoro, M. M., and Bolen, D. W. (1992) A test of the linear extrapolation of unfolding free energy changes over an extended denaturant concentration range. Biochemistry. 31, 4901–4907

61. Vranken, W. F., Boucher, W., Stevens, T. J., Fogh, R. H., Pajon, A., Llinas, M., Ulrich, E. L., Markley, J. L., Ionides, J., and Laue, E. D. (2005) The CCPN data model for NMR spectroscopy: development of a software pipeline. Proteins. 59, 687–696

62. Skinner, S. P., Fogh, R. H., Boucher, W., Ragan, T. J., Mureddu, L. G., and Vuister, G. W. (2016) CcpNmr AnalysisAssign: a flexible platform for integrated NMR analysis. J. Biomol. NMR. 66, 111–124

63. Maciejewski, M. W., Schuyler, A. D., Gryk, M. R., Moraru, I. I., Romero, P. R., Ulrich, E. L., Eghbalnia, H. R., Livny, M., Delaglio, F., and Hoch, J. C. (2017) Nmrbox: A resource for biomolecular NMR computation. Biophys. J. 112, 1529–1534

64. Harwig, M. C., Viana, M. P., Egner, J. M., Harwig, J. J., Widlansky, M. E., Rafelski, S. M., and Hill, R. B. (2018) Methods for imaging mammalian mitochondrial morphology: A prospective on MitoGraph. Anal. Biochem. 552, 81–99

